# Integrating coastal microbiome observations for human, oyster and environmental protection

**DOI:** 10.64898/2026.03.02.708974

**Authors:** Raffaele Siano, Isabelle Arzul, Nicholas Chomérat, Angelique Gobet, Cyril Noël, Enora Briand, Mathieu Chevalier, Germain Chevignon, Anaïs Crottier, Patrick Durand, Christine Félix, Sylvaine Françoise, Camille Gianaroli, Tania Hernández-Fariñas, Luc LeBrun, Cyrielle Lecadet, Soizick Le Guyader, Laura Leroi, Charlotte Mary, Chloé Mason, Sylvain Parnaudeau, Jean-François Pépin, Jean-Côme Piquet, Julien Quéré, Sophie Schmitt, Joelle Serghine, Jean-Luc Seugnet, Ophelie Serais, Aouregan Terre-Terrillon, Michèle Gourmelon

**Affiliations:** Ifremer, DYNECO, F-29280 Plouzané, France; Ifremer, ASIM, Adaptation et Santé des Invertébrés Marins, La Tremblade, France; Ifremer, COAST, F-29900 Concarneau, France; MARBEC, Univ Montpellier, CNRS, Ifremer, IRD, Sète, France; IFREMER, IRSI – Service de Bioinformatique (SeBiMER), Plouzané, France; Ifremer, PHYTOX, GENALG, Nantes, France; Ifremer, ODE/COAST/LERN, Port en Bessin, France; Ifremer, LSEM/MASAE/RBE, Nantes, France; IFREMER, ODE/COAST/LERPC, 17390 La Tremblade, France

**Author notes:** co-first authors (Raffaele Siano, Isabelle Arzul, Nicholas Chomérat, Angelique Gobet, Cyril Noël).

**Keywords:** coastal observatory, coastal ecology, microbiology, pathogens, oysters, coastal management

## Abstract

The *Réseau d’Observatoires de Microbiologie Environnementale intégrée* (ROME) was a pilot study conducted in France from September 2020 to August 2023 aiming to establish a network of eDNA-based observatories across four estuarine ecosystems associated with oyster farming: the Bay of Veys (Normandy), the Bay of Brest (Brittany), Marennes-Oléron (Nouvelle-Aquitaine), and the Thau Lagoon (Occitania). Within a One Health framework, the study assessed the influence of river inputs on estuarine microbiome structuring and the emergence of microbiological hazards affecting human, aquaculture, and ecosystem health.

Over 2,000 samples were collected during the study, including biweekly surface water and monthly adult oyster samples. Environmental nucleic acids were analysed using metabarcoding (bacterial and protist communities) and metagenomics (human RNA viruses). The coastal microbiome, including pathogenic and harmful taxa relevant to humans and aquatic invertebrates, was characterized. River influence on microbial community composition was examined through spatial comparisons of stations exposed to varying levels of freshwater runoff, while oysters acted as bio-integrators of local microbial diversity.

Results revealed coherent coastal-to-offshore microbiome structuring across all ecosystems, with local variations linked to riverine inputs. eDNA metabarcoding allowed to detect a wide range of prokaryotic and eukaryotic pathogens, as well as harmful algal bloom (HAB) genera, several not captured by conventional monitoring. These findings demonstrate the potential of the ROME eDNA observatory network for high-resolution, integrative surveillance of microbial biodiversity and early detection of biological risks in estuarine environments.

## Introduction

Coasts are the most densely populated areas of the planet with a population of about 2.4 billion people (about 40% of the world’s population) living within 100 km of the coast (Burke, 2001). Marine coastal areas are crucial to the Earth’s system, serving as the interface between land and sea. Their biodiversity plays an important role in providing valuable ecosystem services, for human health and for the global economy. The biogeochemical processes occurring at this interface support the global ocean system and sustain a wide variety of organisms. This land-sea interface is particularly significant in estuarine ecosystem functioning, where freshwater from land drainage mixes with seawater (Pritchard, 1989) establishing a hydrological connectivity between the land and sea (Bracken et al., 2013). River run-offs are the primary drivers of estuarine ecosystem functioning, and their impact can vary due to human activities (e.g., agriculture, urbanization, industrial pollution), seasonal natural fluctuations (e.g., rainfall), and extreme climatic events (e.g., storms, floods, droughts). The increasing intensity and frequency of these events makes the study of estuarine spatiotemporal dynamics more complex. Furthermore, human activities such as aquaculture, marine transport, harbour operations, and tourism contribute to organic and inorganic matter and pollutant inputs that can affect biogeochemical cycles and biodiversity (Martínez et al., 2007; He and Silliman, 2019).

River run-offs create gradients of organic, inorganic matter and salinity which are key drivers of microbial community composition, shifts, and functioning (Smith et al., 2010; Fortunato et al., 2012; They et al., 2015; Urvoy et al., 2022). Estuarine microbiomes are specific, highly variable communities of prokaryotic and microeukaryotic organisms (Logares et al., 2009; Telesh and Khlebovich, 2010) resulting from a mix of freshwater species from land, brackish halotolerant species, resuspended benthic species, and diluted marine species. In addition, anthropic pressures may lead to the emergence of biological risks such as microorganisms potentially pathogenic to humans: faecal bacteria (pathogenic *Escherichia coli*), zoonotic bacteria (*Salmonella* spp. and *Campylobacter* spp.) and enteric viruses (Leight et al., 2018; Rincé et al., 2018; Desdouits et al., 2023), blooms of harmful algae producing toxins which have strong negative impacts on animal and human health (Lassus et al., 2016), and pathogenic microorganisms on farmed shellfishes (Foster et al., 2011; Moreau et al., 2015). The impact of fluctuating river run-off inputs on these microbial communities—both in terms of quantity and composition—is not fully understood yet. In a context of global climate change and increasing anthropic pressures, microbial communities are of paramount importance, and therefore the study of their composition, dynamics and functioning requires comprehensive approaches that consider all the natural and anthropogenic factors for the development of less destructive human practices (Nogales et al., 2011).

Genomic techniques, particularly environmental DNA (eDNA) analysis (as defined by Ficetola et al., 2008; Taberlet et al., 2018; Pawlowski et al., 2020) provide enhanced insight into microbiome diversity and function (Zinger et al., 2012). In particular, biodiversity surveys based on amplicon sequencing (*i.e.* metabarcoding) of nucleic acids in the environment and associated with a host have been widely used for ecological studies, biomonitoring, and ecosystem management (Berry et al., 2021). While metabarcoding still faces challenges such as DNA decay and incomplete reference databases, this approach has yielded significant discoveries, including the identification of large-scale biogeographic patterns (de Vargas et al., 2015; Ibarbalz et al., 2019; Logares et al., 2020), the uncovering of biotic interactions (Lima-Mendez et al., 2015), and a better understanding of the rare biosphere (Sogin et al., 2006; Gobet et al., 2012; Pawlowski et al., 2020).

Beyond microbial community exploration, eDNA metabarcoding is also of main interest for environmental management. Traditional microbial biomonitoring methods (e.g., microscopy, strain culturing, biological bioassays) are increasingly complemented by eDNA-based approaches. These methods now serve as tools for ecosystem health assessments, biomonitoring, conservation efforts, and bioindicator development (Aylagas et al., 2014, 2016; Cordier et al., 2021; Pawlowski et al., 2021). However, effective implementation of eDNA metabarcoding for biomonitoring requires systems aligned with environmental impact assessment and aquatic resource protection goals. Long-term eDNA observations or targeted spatio-temporal surveys in both impacted and protected sites can reveal shifts in community composition or the emergence of rare species (e.g., pathogens). These shifts could be used as proxies for the conservation of environmental health and the sustainability of exploited resources. eDNA analysis is thus promising to complement data from classical monitored parameters and to provide new tools for stakeholders in coastal marine areas.

Traditionally, monitoring systems analysing the quality of the environment and biomonitoring surveys targeting resource protection and sustainability are carried out independently (Granit et al., 2017). However, it is now recognized that the protection of exploited resources (e.g., aquaculture), particularly in terms of safeguarding human health through consumption, cannot be separated from the preservation of environmental quality (as described by the One Health concept (Wildlife Conservation Society, 2004). Still, few national marine monitoring systems worldwide have adopted this holistic perspective (Buttigieg et al., 2018) especially when resource safety and environmental status are threatened by pollutants. Marine microbiome coastal observatories tend to focus either on the protection and sustainability of exploited resources or on human health (Buttigieg et al., 2018). Environmental surveys incorporating eDNA observations are also rarely integrated with resource or public health monitoring. Most microbial eDNA observations have been initiated by local laboratories in the past 10-20 years to complement traditional biodiversity observatories (e.g., microscopy data), providing new insights into method development and offering foundational data for specific biological and ecological research (e.g. SOMLIT-ASTAN, Caracciolo et al., 2022). However, the establishment of long-term eDNA observatories for biomonitoring and coastal management at the national level is still under development (e.g., the Marine Microbiome initiative by IMOS in Australia, Australian Microbiome Consortium), and more national initiatives are needed.

To integrate and structure coastal microbial observations for the protection of human health, aquaculture, and the environment, a three-year national pilot study (ROME: *Réseau d’Observatoires de Microbiologie Environnementale Intégrée - Integrated Environmental Microbiology Observatory Network*) was carried out across French coastal areas. Samples from both water and oyster tissues were collected and analysed in a standardized manner across four coastal ecosystems that include economically important oyster farms and are affected by river run-off. By using metabarcoding and RNA-metagenomics, this study aimed to evaluate: (i) the inshore-offshore variability of coastal microbial diversity (bacteria and protists) in areas impacted by river run-off and oyster breeding, (ii) the similarities in community structure and composition between water samples and oyster tissues, and (iii) the emergence of potential pathogens (human viruses, animal or human pathogenic bacteria, harmful photosynthetic microorganisms (phytoplankton and cyanobacteria), oyster parasites) in water and oyster tissues. To our knowledge, this is the first attempt in Europe to create an integrated national microbial observatory system that not only addresses academic questions but also provides information to develop new tools for stakeholders to manage coastal environments and associated exploited resources. This work presents the ROME project and its associated metabarcoding and metagenomic databases. It provides global diversity analyses and conclusions aimed at developing an integrated eDNA and oyster microbiome monitoring network at a national level.

## Material and Methods

### Sampling strategy and study sites

Estuarine microbiomes were monitored for three years (September 2020 - August 2023) at four ecosystems of the French metropolitan coast located, from North to South, in the English Channel, Atlantic Ocean and Mediterranean area. These coastal ecosystems, namely Bay of Veys (BV), Bay of Brest (RB), Marennes-Oléron (MO), and Thau lagoon (ET) encompass different bioclimatic zones, are characterized by markedly different hydrological conditions and tidal regimes and include four of the most important oyster farming areas of the country.

In each coastal ecosystem, water was collected fortnightly at two sampling sites: off-shore (OS) and in-shore (IS), the latter being more influenced by river run-offs and anthropic activities including oyster farming. In addition, oysters (*Magallana gigas*) were sampled (OY) monthly nearby OS (BV-OY and ET-OY) or IS (RB-OY and ET-OY) sites (Appendix Table 1). At the BV, RB, and MO coastal sites, oysters were collected from intertidal areas where low tides expose them to air, whereas at the ET site, oysters remain permanently submerged. Oysters from BV, RB, and ET sites were harvested in category B shellfish production areas (*i.e.* where 90% of samples must contain ≤ 4,600 *Escherichia coli (E. coli,* proxy of faecal bacteria contamination) per 100g shellfish flesh and all samples must contain < 46,000 *E. coli*/100g; .EU Regulation 2019/627). Category B shellfish must undergo purification or relaying before being marketed for human consumption according to this EU regulation. Over the three years of sampling a total number of 584 water (IS + OS) and 161 oyster samples were collected. At each station, temperature and salinity were measured *in situ* using several portable probes: Ysi pro 30 (BV), WTW cond 3210 (RB), Ysi Exo 2 (MO), and WTW multi 3510 ids with a tetracon 925 sensor (ET).

The Bay of Veys (BV) is located in the English Channel, West to the Seine River Bay, and constitutes an intertidal estuary area of 37 km^2^ receiving four main rivers (Vire, Aure, Douve, and Taute). This area is economically important for all shellfish aquaculture (about 10,000 tons per year), particularly oyster production. The OS station was located in the eastern lateral part of the bay, under the influence of the four rivers, and it has been monitored for water quality, nutrients, and phytoplankton in the frame of the French REPHY monitoring system since 1984. The IS station was located well upstream at the confluence of Aure and Vire rivers near Isigny-sur-Mer. Oysters were collected in the eastern side of the Bay, close to the OS station (Fig. 1)

**Figure 1.**
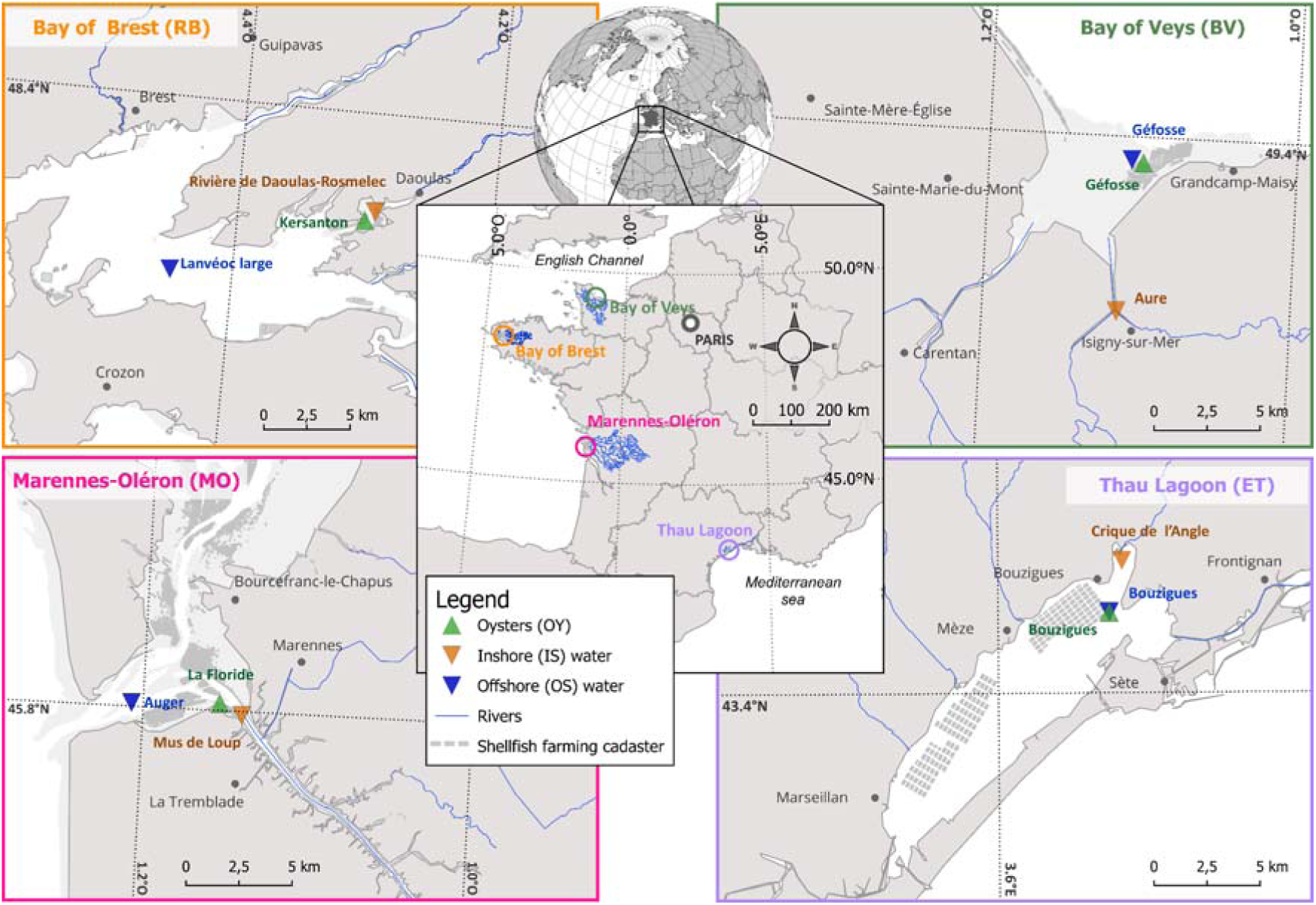
Localization of the three sampling sites (oysters (OY), inshore (IS) and offshore (OS) waters) for the four studied coastal ecosystems (Bay of Brest (RB), Bay of Veys (BV), Marennes-Oléron (MO) and Thau lagoon( ET))

The Bay of Brest (RB) is a large bay of 180 km^2^ of the Atlantic Ocean and characterized by a strongly jagged coastline (∼ 350 linear km) highly influenced by the tidal cycle. The OS station was located in the middle of the bay, where waters from the open ocean (western part) and those from large estuaries of Elorn and Aulne rivers (eastern part) mix. This station has been sampled for water quality (in the context of the Water Framework Directive) and phytoplankton since 1993 in the frame of the French monitoring system REPHY. The IS was located in the eastern part of the bay, and is under the influence of a short coastal river (Mignonne, 21 km long) with a drainage basin of 111 km^2^. Although the corresponding watershed is relatively sparsely populated, with almost no industrial activity, it is strongly impacted by cattle and pig livestock farming, and agriculture (Mauffret et al., 2012). The OY station was close to the IS station.

The Marennes-Oléron (MO) ecosystem is located in the Atlantic Ocean, and includes the most important shellfish production of France (> 50,000 tons per year, mostly oysters) (Agreste, 2018, 2021, 2024). It has also an essential role in spat collection for oysters and mussels. This area is characterized by a large tidal range (> 6 m) and strong currents (3m/s), as well by the influence of the Gironde River from the south. The very high tidal range generates powerful currents at high tide that can overwhelm riverine inputs, leading to a reversal of the usual hydrodynamic regime. The geomorphological configuration of the Gironde and Seudre estuaries, which are long, shallow, and have a gentle slope, favours the upstream penetration of seawater, which enhances the reversal dynamic, particularly during spring tides (Latouche and Jouanneau, 1990). The OS station was located in the Maumusson strait, south to Oléron Island and has been monitored for phytoplankton and water quality (in the context of the Water Framework Directive) since 1984 and 2006, respectively (Ratiskol, 1994; Pellouin-Grouhel and Romana, 2006). The IS station was located in the estuary of Seudre River, close to La Tremblade city and at the outlet of the wastewater treatment plant. OY station was located between IS and OS, in correspondence with the foreshore zone of oysters and clams production.

The Thau lagoon (ET) is one of the largest Mediterranean lagoons (68 km^2^), with a large drainage basin (280 km^2^) widely occupied by natural, agricultural, and urban areas (essentially in the Northern part, Fiandrino et al., 2003, 2017). The lagoon is euhaline, shallow (4.5 m in average), with no tide, and is mostly used for fishing and aquaculture (production of mussel and oysters, reaching 10% of national oysters production). The Thau lagoon is economically important for the Occitanie region as shellfish farming production represents around 8-10,000 t.yr^−1^ since 2010 (Derolez et al., 2020a). The Thau lagoon experienced eutrophication in the 70’-80’, due to large nutrient loadings from its watershed at that time (Souchu et al., 1998; La Jeunesse and Elliott, 2004; Plus et al., 2006). The installation of wastewater treatment plants on the watershed led to a drastic reduction of nutrient inputs to the lagoon, largely lowering its eutrophication status, making the lagoon ecologically “healthy” according to the EU Water Framework Directive (Derolez et al., 2019; Pete et al., 2020). However, the reduction in nutrient input led to a phytoplankton community shift and biomass reduction, which may threaten the shellfish industry (Derolez et al., 2020b). The OS station was located in the south-eastern part of the Bouzigues production area and has been monitored for water quality (nutrients since 2016, chemical contaminants since 1979), phytoplankton (since 1987) and microbiology (since 1988). The IS station was located in the northeastern corner of the lagoon (*Crique de l’Angle*) under the influence of the river Vène. The river is the most important permanent freshwater alimentation of the lagoon and receives wastewater from treatment plants of Montbazin. The OY station was located close to the IS site and the Vène river outlet.

### Water sample analyses Water Filtration

Subsurface water (-1 m) was collected using a Niskin bottle. For ecosystems influenced by tides (i.e. BV, RB, and MO coastal sites), water was sampled ± 2 h during high tide at OS, and during lowering tide at IS. Once collected, water was stored in opaque containers for transportation to local Ifremer laboratories. Samples were kept refrigerated and filtered within 24 h.

To sort micro-plankton (> 20 µm), micro-nano-plankton (> 3.0 µm) and pico-plankton (0.2 – 3.0 µm), 4 liters of water were filtered using a 20 µm filter (GVS – 1215078, diam. 47mm) and 3 liters were filtered serially onto 3 µm (Millipore GTTP14250, 142mm) and 0.2 µm (Millipore TSTP14250, 142 mm) polycarbonate filters. No prefiltration of water was carried out before sample collection. Filtration was carried out by a peristaltic pump (Minipuls3, Gilson) at a speed of 44 rpm. When filters clogged, another filter was used to systematically collect the same water volume. At each sampling date, filtration systems were systematically cleaned with milliQ water and bleach filtered onto the 20 µm and the 3-0.2 µm filtration systems between two samples (IS and OS) of the same series. After filtration, each filter was cut in two equal parts using a sterile scalpel blade. Half filters were rolled, placed in cryotubes, flash-frozen in liquid nitrogen and stored at -70 °C until DNA extraction. DNA was extracted from one half of the filter. The second half-filter was stored at -70 °C for biobanking. A total number of 1752 filters were obtained.

In parallel, 1 liter of subsurface water was analysed for viral fraction using a combination of two concentration methods: a filtration-based protocol developed for sewage (Katayama et al., 2002) and a flocculation-based protocol designed for seawater (John et al., 2011; Alberti et al., 2017). Briefly, water samples were directly filtered on a negatively-charged HA-type membrane with 0.45 μm pores (Millipore, Burlington, MA, USA) placed on a vacuum sterile bottle. Filtrate was kept, and filters were rinsed with 100 mL of 0.5 mM H2SO4 (pH 3) prior to viral elution with 1 mM NaOH (pH 10.5). After pH neutralization, 2 mL of viral concentrate was obtained. In parallel, 200 μL of 10 g/L FeCL3 solution was added to the filtrate kept at 4 °C, and incubated 2 h at 10 °C under gentle agitation, in the dark. A flocculate was then collected on a 0.8 μm pore-size polycarbonate filter (Whatman, Maidstone, UK). Virus resuspension was achieved with 2 mL of ascorbate-oxalate–EDTA buffer during a 30 min incubation at 4 °C under agitation.

### Water nucleic acid extractions

Planktonic DNA was extracted using the NucleoSpin Plant II Mini Kit for micro-nano-plankton and pico-plankton (Macherey-Nagel, Düren, Germany, Ref. 740770. 50) and the NucleoSpin Plant II Midi Kit for micro-plankton (> 20 µm) (Macherey-Nagel, Düren, Germany, Ref. 740771. 20), respectively following manufacturer instructions. Prior to extraction, an additional lysis step was performed at 56 °C for 2 h with Proteinase K (Macherey-Nagel Ref. 740506) and lysozyme (Sigma-Aldrich Ref. 4403-5g, Saint-Louis, MO). Finally, DNAs were eluted in 100 µL (NucleoSpin Plant II Mini Kit) or 200 µL (NucleoSpin Plant II Midi Kit) PE (2 min., 65°C). One DNA extraction blank was performed for each new kit used. DNA concentration was measured using the picogreen 2020-2021 (Quant-iT Picogreen Kit (Invitrogen, Carlsbad, Californie) and AccuBlue® High Sensitivity dsDNA Quantitation Kit (Biotium, Fremont, CA) and adjusted to 5 ng.µL^−1^. Aliquots of each sample were biobanked (Labcollector) and stored at -70 °C.

The 4 mL of viral suspensions obtained were extracted using the NucliSens kit (bioMérieux, Lyon, France) with 10 mL of lysis buffer and 140 L of magnetic silica, and eluted in 100 μL of the kit’s elution buffer (Desdouits et al., 2021).

### Water PCR amplifications

Bacterial and protist planktonic diversity were analysed by amplifying the V4-V5 variable region of the 16S rDNA and V4 variable region of the 18S rDNA, respectively. DNA amplification was done by PCR, in triplicates. V4-V5 16S rDNA was targeted using the primers 515F-Y (5’ GT GYC AGC MGC CGC GGT AA 3’) and 926R (5’ CC GYC AAT TYM TTT RAG TTT 3’, Parada et al., 2016). V4 18S rDNA was targeted using the following set of primers : TAReuk454FWD1 (5’ CC AGC ASC YGC GGT AAT TCC 3’) and TAReukREV3 (5’ AC TTT CGT TCT TGA TYR A 3’, Stoeck et al., 2010). Each primer contains the Illumina sequencing adapter (from Genotoul platform). For V4-V5 16S rDNA amplification, the PCR mix was composed of 0.12 µL 5U/µL GoTaqFlexi G2 DNA Polymerase (Promega), 0.6 µL (10 mM) dNTP, 0.45 µL (20 μM) forward primer, 0.45 µL (20 µM) reverse primer, 18.58 µL molecular-grade water, 6 µL (5X) buffer, 1.8 µL (25mM) MgCl2, and 2 µL of extracted DNA. The PCR program included an initial denaturation step at 95 °C for 5 minutes followed by 32 cycles at: 95 °C (30 s), 50 °C (60 s), 72 °C (60 s). For V4 18S rDNA amplification, the PCR mix was composed of 0.3 µL 2U/µL Phusion High-Fidelity DNA Polymerase (Thermo Scientific), 0.6 µL (10 mM) dNTP, 0.9 µL (3% final) DMSO, 1.1 µL (10 μM) forward primer, 1.1 µL (10 µM) reverse primer, 18 µL molecular-grade water water, 6 µL (5X) buffer and 2 µL of extracted DNA. The PCR program consisted of an initial denaturation step at 98 °C for 30 s, followed by two sets of cycles (1) 12 cycles at: 98 °C (10 s), 53 °C (30 s), 72 °C (30 s), and (2) 18 cycles at: 98 °C (10 s), 48 °C (30 s), 72 °C (30 s). For both markers, the last step was a final extension at 72 °C for 10 minutes. Electrophoresis gels were used to check the amplification. In case of no amplification on the first gel, PCR was repeated from a higher dilution of the DNA extracts. Sample triplicates were pooled for a final volume of 75 µL.

### Oyster microbiome sample analyses Oyster sampling

For oysters, 15 adult oysters (*Magallana gigas)* were collected monthly and kept refrigerated upon transportation to the laboratory. Oysters were then processed as follows. 1st batch: ten oysters were opened and the digestive tissues (hereafter described as DT), where the microorganisms brought by the oyster filtration are concentrated, were removed by dissection. After fine cutting and homogenization, 2 g of DT from these 10 specimens were stored at -70 °C for metagenomic analysis of human RNA viruses. 2nd batch: the remaining five oysters were opened and all tissues were collected, cut and homogenized. A sample (approx. 7 g) of these homogenized tissues were “flash-frozen” in liquid nitrogen and then ground. The resulting powder was stored at -70 °C for metabarcoding analysis (Supp Fig. 1).

### Oyster Nucleic Acid extraction

To investigate bacteria and protist diversity from oysters in a simultaneous manner, nucleic acids were extracted from ground tissues from the 2nd batch following Strittmatter (2019). DNA and RNA were extracted simultaneously using the ALLPREP mini kit kit (Qiagen ref. 80204, Hilden, Germany). DNA was used for 16S rDNA and RNA for 18S rDNA amplification, after retro transcription. Nucleic acid elution was done in 100 µL of sterile molecular grade water (RNA) or elution buffer (DNA). DNA and RNA were stored at -70 °C until PCR or RT, respectively. DNA concentration was measured using the picogreen 2020-2021 Quant-iT Picogreen Kit (Invitrogen, Carlsbad, Californie). RNA concentration was measured with an EPOCH spectrophotometer.

To investigate human RNA viruses, nucleic acid extraction was carried out from DT homogenates (Strubbia et al., 2020). Briefly DT aliquots (2 g) were lysed with proteinase K (30 U.mg^−1^, Sigma-Aldricht, St-Quentin France), sonicated before centrifugation (5 min 3,000 g). Supernatant was mixed with sodium pyrophosphate (10 mM final concentration) and incubated for 40 min at 4 °C under gentle agitation. After centrifugation for 20 min at 8000×*g*, supernatant (approximately 3 mL) was recovered, mixed with 1.5 mL of polyethylene glycol 6000 (PEG 24% wt/vol, Sigma-Aldrich)-sodium chloride (1.2 M), and incubated for 1 h at 4 °C. After centrifugation for 20 min at 10,000×*g*, the pellet was resuspended in 2 mL of preheated (56 °C) glycine buffer (0.05 M; pH 9), then filtered using 5, 1.5, and 0.45 μm acetate cellulose filters. Filtrate was treated with 20 μL of OmniCleave™ Endonuclease (Lucigen corporation; 200 U/μl) and 200 μL of MgCl_2_ (100 mM) for 1 h at 37 °C. The extraction of nucleic acids was carried out using the NucliSENS kit (bioMerieux) with the semi-automatized eGENE-UP™ system (bioMerieux). Nucleic acids were treated by TURBO DNase (25 U) for 30 min at 37 °C (Ambion, Thermo Fisher Scientific, France). An additional RNA purification was carried out using the RNA Clean - Concentrator™-5 kit (Zymo Research, Irvine, United States). Nucleic acids were recovered in 100 μL of the elution buffer and stored at −70 °C.

### Oyster PCR amplifications for bacteria and protists

Oyster microbiome diversity in oyster sample was analysed from DNA and cDNA by amplifying the V3-V4 region of the 16S rDNA using the primers Bakt_341F (5’ CCT ACG GGN GGC WGC AG 3’) and Bakt_805R (5’ GAC TAC HVG GGT ATC TAA TCC 3’, Klindworth et al., 2013) and the V1-V2 region of the 18S rDNA using the primers PCR1F (5’ ACC TGG TTG ATC CTG CCA 3’) and PCR1R (5’ GTA RKC CWM TAY MYT ACC 3’, Clerissi et al., 2018), respectively. For protists, a plate of extracted RNA from ground oyster tissue (8 µL RNA/well) was prepared following quantification of total RNA (EPOCH spectro assay). RNA extracted from oysters was treated with DNAse (DNase I, ref AMPD-1KT, Sigma) and retro transcripted in cDNA using the Iscript cDNA Synthesis kit (Bio-Rad, ref 1708891, Hercules, CA). For bacteria, amplifications were carried out in a total volume of 30 μL using 2 µl of DNA, 6 μL (5X) Buffer HF, 0.6 µL (10 mM) dNTP, 0.45 μl (20 μM) of each primer, 0.3 µL of 2U µL-1 Taq Phusion High-Fidelity DNA Polymerase (Thermo Scientific) and 20.2 µL of molecular grade water. PCR1 conditions were: 5 min at 98 °C, followed by 30 cycles of 10 s at 98 °C, 30 s at 55 °C, and 20 s at 72 °C, and final elongation for 5 min at 72 °C. For protists, amplifications were carried out in a total volume of 30 μL using 5 µL of cDNA, 6 μL (5X) Buffer Go Taq, 1.8 µL (25 mM) MgCl2, 1.2 µL (10 mM) dNTP, 0.48 μL (20 µM) of primer PCR1F_Clerissi, 2.4 μL (20 μM) of primer PCR1R_Clerissi, 0.2 µL of 5 U µL-1 GoTaqFlexi G2 DNA Polymerase (Promega) and 12.92 µL of molecular-grade water. PCR1 conditions were: 5 min at 95 °C, followed by 35 cycles of 60 s at 95 °C, 30 s at 52 °C, and 30 s at 72 °C, and final elongation for 7 min at 72 °C. PCR products were loaded onto an agarose gel to ensure correct amplification. If no amplification was observed, PCR was repeated from a higher dilution of the DNA extracts. Aliquots of each sample were biobanked (Labcollector) and stored at -70 °C.

### Oyster metagenomics analyses for human RNA viruses

Metagenomic analysis of RNA viruses in the first-year samples required two runs (NovaSeq, Illumina) with the same number of samples and negative controls. cDNA reverse-transcription of RNA extracts was performed using the enzyme Superscript IV (Thermofisher, France) and random hexamer primers (Thermofisher, France). After the production of cDNA using the second strand reaction buffer and synthesis enzyme mix (NEBNext Ultra RNA Library prep, New England Biolabs, France), a physical fragmentation (Ultrasonicator M220, Covaris) was carried out for 110 s (Strubbia et al., 2020). Libraries were prepared using the KAPA Prep Kit (Roche, France). After ligation of the adapters, several libraries cleanup steps using AmPure XP Beads (Beckman Coulter, United States), and 80% ethanol were carried out to select fragments between 150 and 500 bp. The libraries were then quantified and pooled equimolarly into seven pools. The sequencing was carried out on the Illumina NovaSeq 6000 using NovaSeq reagent Kit to generate 2×250 base pair reads at the ICM platform (Institut du Cerveau, Paris, France). Two runs with the same number of samples and negative controls were performed.

### Nucleic acid sequencing

For each sample of water and oyster, PCR products from the three replicates were pooled and 50 µL of the suspension were sent to the GeT-PlaGe platform (Toulouse, France), which performed: (i) purification and quantification of PCR amplicons; (ii) amplification (12 cycles) using Illumina-tailed primers allowing the addition of a dual-index specific for each sample; (iii) preparation of an equimolar mixture of the dual-index amplicons; (iv) Illumina MiSeq sequencing using the V3 (600 cycles) paired-end sequencing kit (250 bp × 2) with an addition of about 30% of PhiX. The quality of the run was checked internally using PhiX, and then each paired-end sequence was assigned to its sample using a demultiplexing script.

### Bioinformatic and biostatical analysis

Raw sequence data were pre-processed using the SAMBA pipeline (v4.0.1) (https://gitlab.ifremer.fr/bioinfo/workflows/samba). SAMBA is a FAIR scalable workflow integrating, into a unique tool, state-of-the-art bioinformatics and statistical methods to conduct reproducible metabarcoding analyses using Nextflow. The workflow is mainly based on QIIME 2 (Bolyen et al., 2019) and DADA2 (Callahan et al., 2016). Briefly, a primer removal step is performed using Cutadapt (Martin, 2011) as well as the removal of reads with incomplete or incorrect primer sequences. The remaining reads are clustered into Amplicon Sequence Variants (ASV) using DADA2 following a four steps approach: quality filtering, sequencing error correction, ASV inference (pairwise read merging) and chimera detection. In order to limit the identification of false positive ASV (PCR biases, remaining sequencing errors), a complementary step of ASV clustering is performed using dbOTU3 (Olesen et al., 2007). The resulting ASVs were filtered based on their taxonomic assignment. The high-quality 16S rDNA ASVs were taxonomically assigned using a Naive Bayesian classifier against the SILVA v138 database (Pruesse et al., 2007), trimmed to the V3–V4 region with primers 341F/805R for water samples and Bakt_341F/Bakt_805R for oyster samples. The 18S rDNA ASVs were assigned using the PR2 database (v5.0.0) (Guillou et al., 2013). For the 16S rDNA data, ASVs classified as Eukaryotes, Archaea, chloroplasts or mitochondria were removed. We also remove the bacterial genus *Ralstonia* that is known as a common contaminant taxa in several studies (Salter et al., 2014; Eisenhofer et al., 2019), and that was found in high sequence abundance in some negative control samples and low microbial biomass samples (e.g. some oyster tissue samples) in our study. From 21512 to 114859 and from 29142 to 124286 sequences per sample were obtained in the water and oyster samples, respectively. Similarly, for the 18S rDNA data, in order to focus on protist communities, ASVs assigned to the following taxa were excluded: Bacteria, Archaea, Streptophyta, Metazoa, Fungi, Florideophyceae, Ulvophyceae and Phaeophyceae. From 29736 to 98483 and from 14009 to 127108 sequences per sample were obtained in the water and oyster samples, respectively.

Statistical analyses were conducted to produce alpha and beta diversity measures using mainly the *phyloseq* (McMurdie and Holmes, 2013) and the *metaBmisc* (https://gitlab.ifremer.fr/cn7ab95/metabmisc) R packages. Briefly, alpha diversity of the microbial communities was assessed using the Shannon diversity index to quantify species richness and evenness within each sample. Beta diversity was evaluated using Bray–Curtis dissimilarities applied on a rarefied count table (rarefaction performed over 1,000 iterations) while differences in community composition between sites were visualised using non-metric multidimensional scaling (nMDS). To identify shared and unique ASVs across the different sampling sites, intersection analyses were performed. To provide statistical support for observed patterns, Permutational Multivariate Analysis of Variance (PERMANOVA) tests were conducted. All figures were generated using the *ggplot2* R package (Wickham, 2016).

For viral metagenomics, we used the pipeline MAEVA (v3.0.0; https://gitlab.ifremer.fr/bioinfo/workflows/MAEVA). This FAIR workflow based on Nextflow tool, starts with cleaning and trimming steps. Clean reads are filtered to remove bacteria RNA reads by mapping to the Silva RNA database (downloaded 2022-03-01) and dereduplicated (CD-hit) to remove PCR duplicates. Then, two different ways of analysis are possible: a direct read mapping on the Virus Pathogen Database (v2.0.2, 10.5281/zenodo.7876308) using Esviritu pipeline (v0.2.3) (Tisza et al., 2023) or de novo assembly using metaSPAdes (v3.14.0, De novo assembly). No contig was assigned as a viral family of health concern (non envelopped RNA viruses). Sequencing quality can be assessed by the number of reads obtained per sample. No site-related effect was observed on the mean number of reads for oyster samples (BV: 4.7M, ET: 5.1M, MO: 5.3M, RB: 4.9M) or for water types. Similarly, both types of water yielded comparable read numbers (IS: 4.1M, OS: 3.9M). Negative controls, prepared from molecular biology–grade water, showed a lower number of reads (1M).

Metadata description was provided for each sampling site and the corresponding data was imported into a specific LabCollector© instance. The Marine Organisms and Resources Storage systEm (MORSE) of Ifremer centralised the information on all biological samples (filters, digestive tissues, oyster ground, DNA and RNA extracts), including a unique reference system for biological samples, information on regulatory obligations (i.e. Access and Benefit-Sharing, ABS).

### Search for pathogen taxa

The pathogen taxa investigated are listed in Appendix Table 2. For bacteria, the genera that include species considered as human and/or animal pathogens (Risk Groups 2 and 3, indicating moderate and high individual risk, respectively) according to Public Health Agency of Canada (ePATHogen – Risk Group Database; https://health.canada.ca/en/epathogen), were selected. This selection was supplemented by other genera reported as pathogenic in previous studies and reviews (Paillard et al., 2004; Austin, 2005; Brenner et al., 2005; Austin and Austin, 2007; Vela et al., 2011; Whitman, 2012; Loch and Faisal, 2015; Travers et al., 2015; Takano et al., 2016; Escobedo-Hinojosa and Pardo-López, 2017) .

For cyanobacteria, the taxonomic reference list of the UNESCO Intergovernmental Oceanographic Commission (https://www.marinespecies.org/hab/index.php) and reports from French national agencies (Afssa and Afsset, 2006; ANSES, 2020) were used to identify potentially toxic taxa.

For Harmful Algal Bloom (HAB) taxa, organisms were sorted into two different groups: (i) taxa potentially toxic to humans (Public Health subsection) and (ii) organisms having potential effect on ecosystems, macrofauna, or animals (Environmental Health subsection). The Taxonomic Reference list from the Intergovernmental Oceanographic Commission of UNESCO (https://data.hais.ioc-unesco.org/) was used to identify HAB taxa with addition of known potentially harmful species/genera in the studied region (such as the dinoflagellate *Lepidodinium*) in the whole dataset. To construct a database including taxonomic ranks as specific as possible and to avoid overlooking some potential harmful taxa, ASVs assigned to the Family rank of Dinophysiaceae, Kareniaceae and Amphidomataceae (Dinophyceae) were also searched in the dataset. A total of 40 genera and 3 Families were selected in the database.

For protist parasites, organisms were sorted depending on their potential impact on humans, bivalves, crustaceans or echinoderms. A total of 17 genera known to include species having a potential impact on public health or marine invertebrate health were searched in the whole datasets.

## Results

### Salinity and temperature conditions across sites

The average salinity observed was different between water types (IS: 26.9 ± 12.7 vs. OS 34.7 ± 2.7) and spatially (BV: 21± 13.7, RB: 31.7 ± 5.4, MO: 32 ± 3.2, and ET: 38.7 ± 1.5). At BV, salinity of the IS waters ranged from 0.2 to 30.8 (mean ± SD: 9.0 ± 8.7), while in the OS waters it varied between 31.7 and 34.3 (33.0 ± 0.5). At RB, IS water salinity ranged from 4.1 to 35.6 (28.7 ± 7.0), compared to 28.8–35.6 (34.1 ± 1.3) for the OS waters. At MO, salinity of IS waters ranged from 17.0 to 35.5 (31.2 ± 3.9), whereas salinity of OS waters ranged from 25.7–35.2 (32.7 ± 2.0). At ET, salinity at IS ranged from 33.8 to 42.0 (38.7 ± 1.5), while salinity at OS ranged from 35.7 to 41.2 (38.7 ± 1.2, Supp Fig. 2A). Temperature was quite homogeneous on average between water types (IS: 15.4 ± 5.7 °C vs. OS 14 ± 4.9 °C) and sampling sites (BV: 14 ± 4.8 °C, RB: 14.6 ± 3.8 °C, MO: 15 ± 4.8 °C, and ET: 16.9 ± 6.8 °C). Water temperature exhibited comparable patterns across sites but with seasonal variations. At BV, the temperature of IS waters ranged from 4.1 to 24.0 °C (14.2 ± 5.2), versus 6.7 to 21.7 °C (13.9 ± 4.4) in the OS waters. At RB, the temperature of IS ranged from 8.5°C to 28.4 °C (15.4 ± 4.5), compared to 9.1–19.2 °C (14.0 ± 3.1) in OS waters. At MO, the temperature of IS waters varied between 6.3 and 23.6 °C (15.1 ± 5.2), while the temperature of OS waters ranged from 6.8 to 22.5 °C (15.0 ± 4.3). At ET, the IS water temperature ranged from 3.9 to 29.9 °C (17.0 ± 7.0), and the OS water temperature from 5.3 to 28.1 °C (16.9 ± 6.7, Supp Fig. 2B).

**Figure 2.**
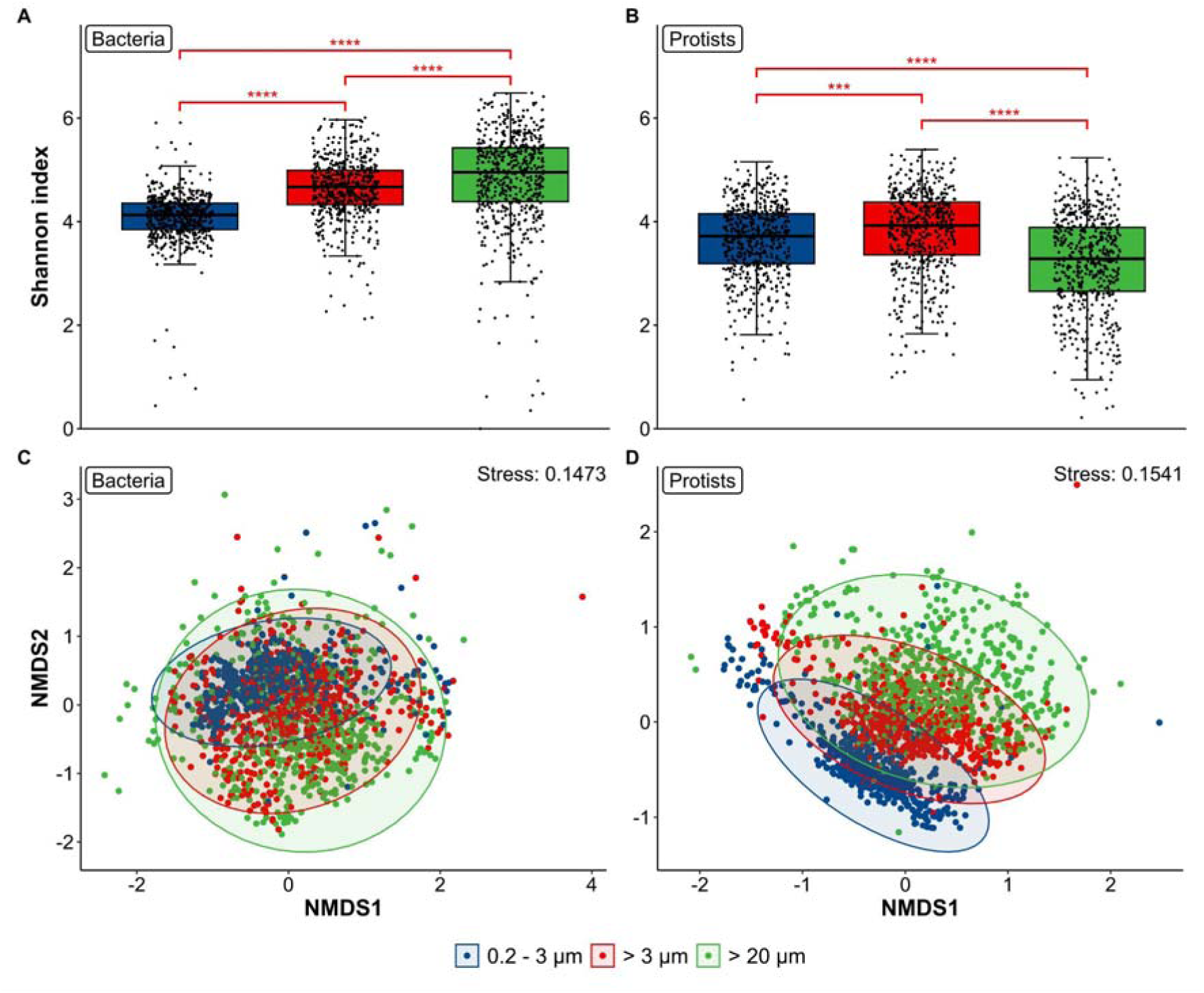
Diversity and structuring patterns of the bacterial (A, C) and protist communities (B, D). Alpha diversity in each sample, calculated using the Shannon index and categorized by size fraction for the bacterial (A) and the protist communities (B). A Kruskal-Wallis test was performed to compare the index distribution across size fractions, followed by a pairwise comparison of the Shannon index for each size fraction using a post-hoc Dunn test with Bonferroni correction for multiple comparisons, as indicated in red above the boxplots. Structure of the bacterial (C) and protist (D) communities in the three size-fractions. Non-Metric Multidimensional Scaling (nMDS) was calculated based on a Bray-Curtis distance matrix. The goodness-of-fit of the two-dimensional representation compared to the original matrix was validated by a low stress value of 14% and 15%, respectively. Samples were coloured according to the size-fraction. Ellipses represent 95% confidence intervals around group centroids. **** p-value < 0.0001, **\*\*\*** p-value < 0.001.

### Bacteria and protist diversity in waters

Microbial diversity varied significantly among the three size fractions (Dunn test: *P* < 0.001, Fig. 2A, 2B). For bacteria, the 0.2–3 µm fraction exhibited a narrower diversity range (Shannon index: 3.8 - 4.3, between 25^th^ and 75^th^ percentiles), compared to the other two size fractions (4.3 - 5 for the >3 µm fraction and 4.4 - 5.4 for the >20 µm). For protists, the diversity distribution was comparable between the two smaller fractions (Shannon index: 3.3 - 4.4) but distinct from the > 20 µm fraction (2.6 - 3.7). When comparing bacterial community structure of bacteria, samples from the three size fractions overlapped (Fig. 2C, PERMANOVA: F = 62.809, R2 = 0.06613, p-value = 0.001, while a clearer sample clustering was observed between size fractions for protists (Fig. 2D, PERMANOVA: F = 88.8432, R2 = 0.06398, *p*-value = 0.001). In both cases, clusters of samples from the larger size fractions overlapped compared to samples clustered according to the 0.2-3 µm, owing to a larger diversity of community structure. For inter-site/station comparisons, only the two size fractions 0.2-3 µm and >3 µm were thus considered to avoid redundant diversity information between the >20 and >3 µm size fractions that were not sequentially separated during water filtering.

Alpha diversity differences among sites were investigated combining data from the three sampling years. One-to-one significant differences between sites were observed for both the bacterial and the protist communities (Fig. 3A, 3B). The interannual alpha diversity variability was analysed combining data of the two selected size fractions and water types. No general trend, but rather a local interannual variability pattern was observed. For instance, for bacteria, the Shannon index distribution differed in RB between Year 1 and Years 2 and 3 (Dunn test: *P* < 0.01) and for protists the third year in MO differed from the 3 other sites (Dunn test: *P* < 0.001, Fig. 3C, 3D). As for the IS vs. OS differences over the three years, bacterial communities significantly differed between RB vs. BV or MO for IS waters and significant differences were detected between MO vs. the three other sites, within OS waters (Dunn test: *P* < 0.01, Fig. 3E). In the protist community across both water types, significant differences were observed between MO and the other three sampling sites (Dunn test: *P* < 0.001, Fig. 3F) and between water types at ET (Dunn test: *P* < 0.001).

**Figure 3.**
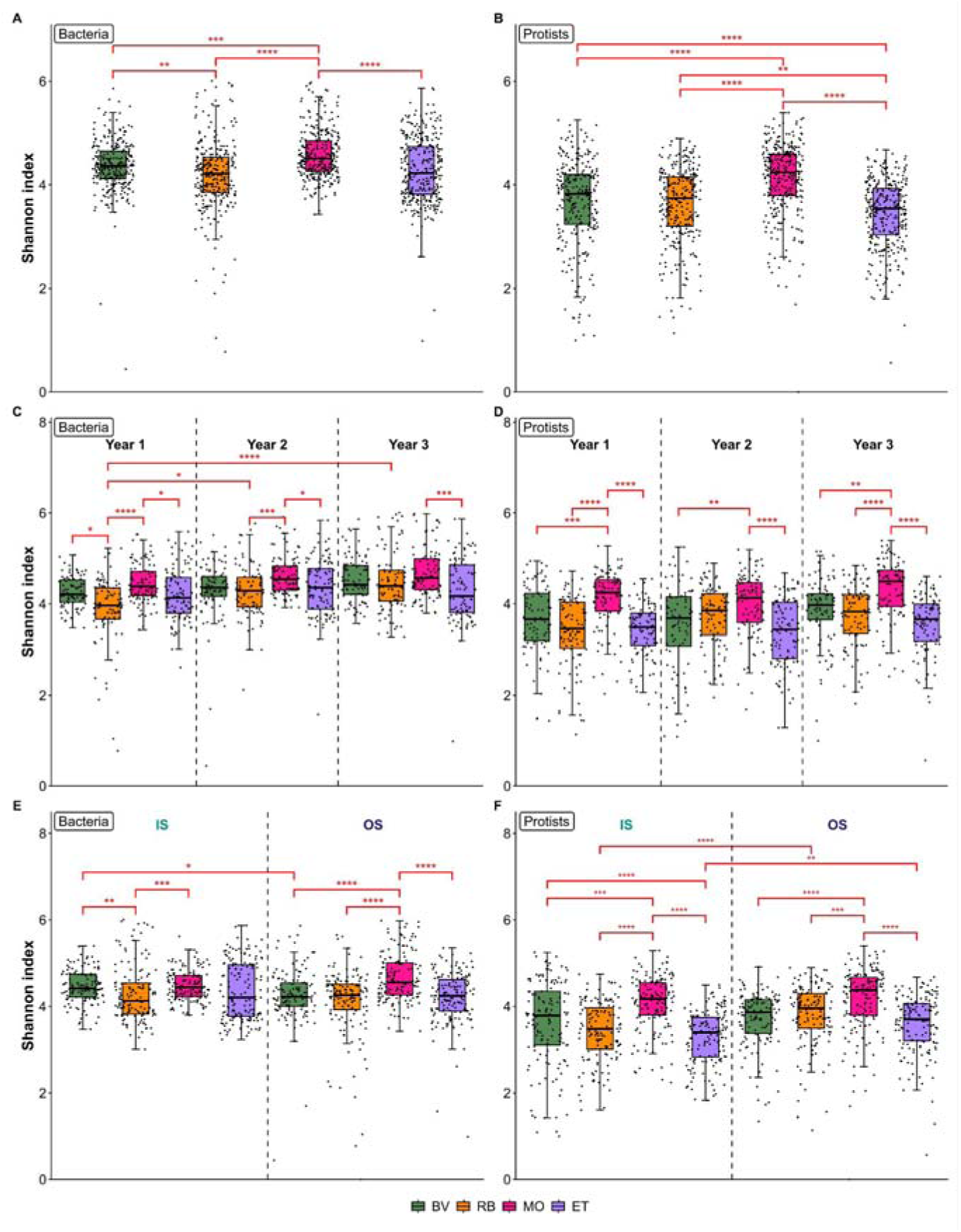
Alpha diversity in each sample of the 0.2-3 µm and the > 3 µm fractions, calculated using the Shannon index for the bacterial (A, C, E) and protist communities (B, D, F), and categorized by sampling site (A, B), by sampling site for each sampling year (C, D), and by sampling site for each water type (E, F). To compare the distribution of the index across sites (A, B), across sampling site for each sampling year (C, D), and across sampling site for each water type (E, F), a Kruskal-Wallis test followed by pairwise comparisons using the post-hoc Dunn test with Bonferroni adjustment for multiple comparisons was performed, as represented in red above the boxplots. BV, Bay of Veys; RB, Bay of Brest; MO, Marennes-Oléron; ET, Thau lagoon. Sampling years were defined as follows: Year 1 corresponds to September 2020–August 2021, Year 2 to September 2021–August 2022, and Year 3 to September 2022–August 2023. IS, inshore waters; OS, offshore waters. **** p-value < 0.0001, **\*\*\*** p-value < 0.001, ** 0.001 < p-value < 0.01 * 0.01 < p-value < 0.05.

### Community structure in waters

Common ASVs to all sites and water types represented only a small percentage of the total ASVs (i.e. 5.2% (985/18,807) for bacteria and 7.7% (930/12,033) for protists), but these accounted for more than 74% of the total sequence abundance for bacteria and protists (Supp. Fig. 3A; Supp. Fig. 3B). By contrast, ASVs specific to site and water type represented in total 53% and 47.7% of the total ASVs for bacteria and protists, respectively, but representing a low sequence abundance (<0.50%) for bacteria and protists (Supp. Fig. 3A; Supp. Fig. 3B). ASV specificity was evident across site, especially in the BV-IS waters (16.3% for bacteria and 18.3% for protists) and RB-IS (10.1% for bacteria and 10.9% for protists), but these ASVs were present in lower sequence abundances (Supp. Fig. 3A; Supp. Fig. 3B). This effect was not seen in waters from the two other sites MO and ET.

The community structure of bacteria and protists in waters was investigated in the four sampling sites. For both microbial groups, samples clustered according to their sampling site. In the Gulf of Biscay, MO samples grouped in a narrower cluster and overlapped with the RB sample cluster (Fig. 4A, Fig.4B).

**Figure 4.**
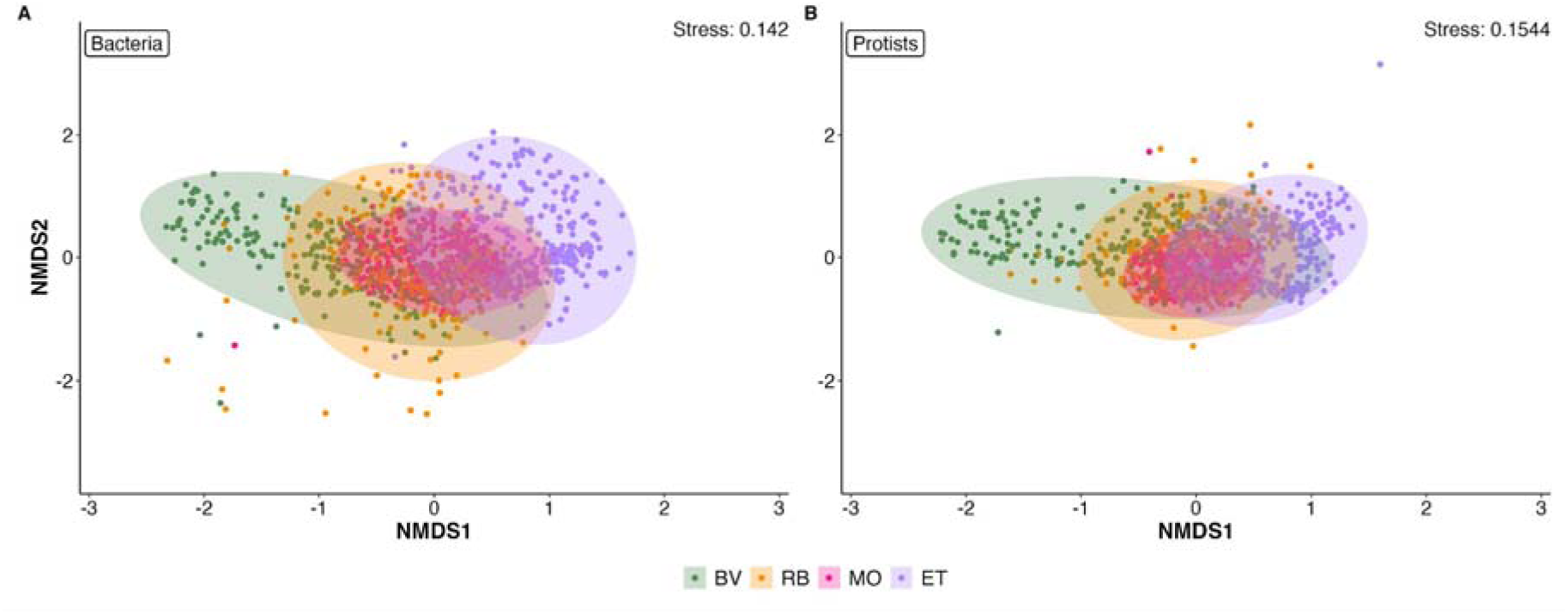
Structure of the bacterial (A) and protist (B) communities in the four sampling sites. Non-Metric Multidimensional Scaling (nMDS) performed on a Bray-Curtis distance matrix. The goodness-of-fit of the two-dimensional representation compared to the original matrix was validated by a low stress value of 14% and 15%, respectively. Samples were coloured according to the sampling sites. Ellipses represent 95% confidence intervals around group centroids. BV, Bay of Veys; RB, Bay of Brest; MO, Marennes-Oléron; ET, Thau lagoon.

Beyond spatial clustering, we further showed that microbial communities clustered according to water type (Bacteria PERMANOVA: F = 53.873, R2 = 0.03452, *p*-value = 0.001; Protist PERMANOVA: F = 22.9880, R2 = 0.01655, *p*-value = 0.001). For all sampling stations, salinity and temperature significantly influenced community structuring with communities in OS sites consistently being associated with higher salinity and temperature (*P* < 0.001, Fig. 5).

**Figure 5.**
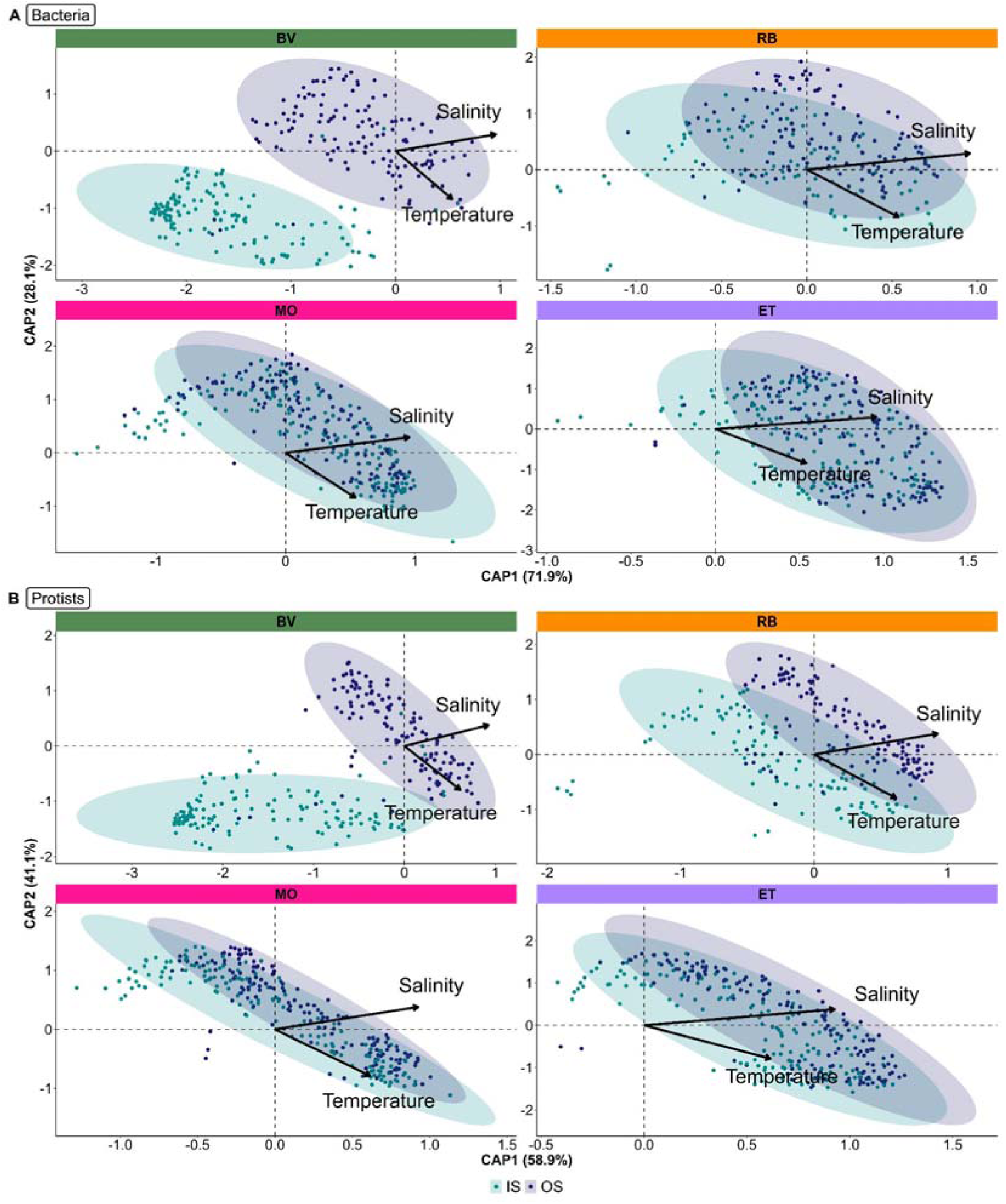
Community structure of the bacterial (A) and protist (B) communities and their relationship with salinity and temperature for each sampling site and in relation to water time. A CAP (Canonical Analysis of Principal Coordinates) was calculated from normalised data. Samples were colored according to water types IS and OS. Ellipses represent 95% confidence intervals around group centroids. BV, Bay of Veys; RB, Bay of Brest; MO, Marennes-Oléron; ET, Thau lagoon.

### Community structure in oyster tissues

Within the oyster batches (OY), a slightly higher proportion of site-specific ASVs was detected compared with the water samples (58.9% in water vs. 60.8% in oysters). These site-specific ASVs were generally of low abundance for both bacteria and protists. Approximately 12.3% (1201/9751) of the bacterial ASVs were common to all sites and accounted for 87% of the relative sequence abundance (Supp. Fig. 3C). For protists, fewer than 10% (236/2534) ASVs were common to all sites, yet they accounted for approximately 62% of the total sequences (Supp. Fig. 3D). By contrast, site-specific ASVs (local biodiversity pattern) represented 60.8% and 53.4% of total ASVs for bacteria and protists, respectively. This pattern was particularly pronounced at BV (18.1% ASVs for bacteria and 16.9% for protists); RB (15.7% for bacteria and 12.7% for protists) and ET (15.6% for bacteria and 14.7% for protists). Despite their high richness, these site-specific ASVs contributed only marginally to overall sequence abundance. For protists, the relative abundance of ASVs common to BV, RB and MO (non-Mediterranean sites) reached 12.7% of the relative abundance with only 5.3% of all protist ASVs, whereas ET-specific ASVs accounted for 2.1% of the relative sequence abundance (Supp. Fig.3D).

For both microbial groups, samples clustered spatially although non-mediterranean samples (i.e. samples from BV, RB and MO) overlapped more than samples from ET, particularly for protists (Fig. 6A, Fig.6B) (PERMANOVA: p-value < 0.001).

**Figure 6.**
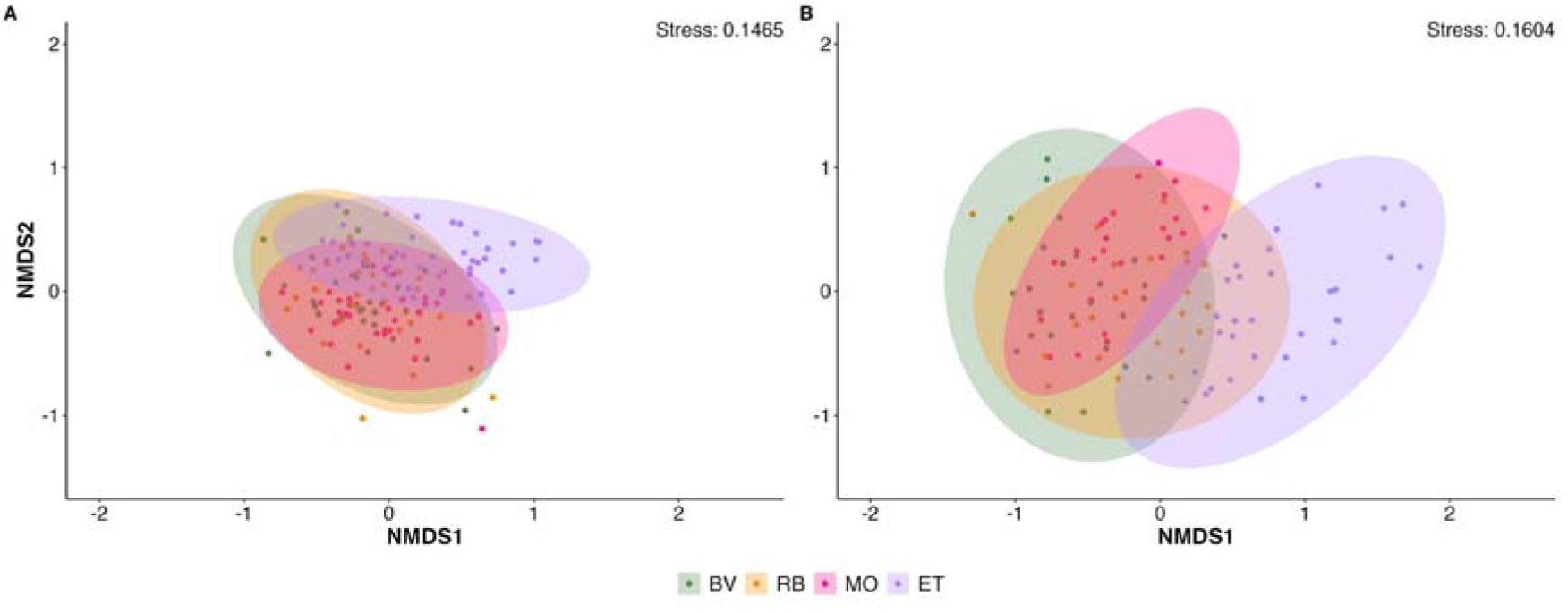
Structure of the bacterial (A) and protist (B) communities in oyster tissues. Non-Metric Multidimensional Scaling (nMDS) performed on a Bray-Curtis distance matrix. The goodness-of-fit of the two-dimensional representation compared to the original matrix was validated by a low stress value of 14% and 16%, respectively. Samples were colored according to the sampling sites. Ellipses represent 95% confidence intervals around group centroids. BV, Bay of Veys; RB, Bay of Brest; MO, Marennes-Oléron; ET, Thau lagoon.

### Taxonomic compositions in waters and oyster tissues

Within the bacteria community, the most dominant orders were Flavobacteriales, Rhodobacterales, SAR11 clade, Pseudomonales, and Burkholderiales, accounting for about 18.9%-38.7%, 7.0%-24.7%, 0.7%-26.9%, 4.3%-16.6%, and 1.4%-24.5% sequences per year, water type and site, respectively (Appendix Table 3A).

At the genus level, in OS waters, SAR11 clade Ia was consistently the most abundant taxon, representing from about 8.1% to 23.1% sequences, depending on the site and year. Within the order Rhodobacterales, the genera *Amylobacter* and *Planktomarina* were particularly abundant in OS waters at BV, RB, and MO, with each genus contributing between about 4.3–6.3% and 3.5–8.7% of sequences across sites, respectively.In contrast, the taxa HBM11 was dominant at ET (3.2-4.2%; Fig. 6A). Among the Flavobacterales order, the NS5 and NS9 marine groups were the most dominant (3.4-6.9% and 1.4-4.6%, respectively). Finally, the most abundant genus within Pseudomonadales was SAR86 (1.8-5.6%). In IS waters at RB, MO, and ET, the genus composition was broadly similar to that of OS waters. The SAR11 Ia clade remained dominant (5.2-18.3%), while the NS3a marine group (Flavobacterales) was notably more abundant in IS compared to OS waters (3.3-8.6% vs. 0.6-1.8% sequences). In contrast, IS waters at BV exhibited a distinct community profile in sequence abundances of *Limnohabitans* (2.6-4.0% vs. 0.00–1.1%) and unknown *Comamonadaceae* (8.9-10.4% vs. 0.01–3.40%; order Burkholderiales*), Flavobacterium* (11.1-13.6% vs. 0.01-3.6%; Flavobacteriales*), Pseudarcicella* (4.7-6.0% vs. 0.01–1.35%; Cytophagales), and the hgcl clade (Frankiales) (3.7-5.7% vs. 0.0-0.7% sequences). Additionally, *Marinobacterium* (Pseudomonadales) was more abundant in IS waters of both BV and RB (1.8-5.0% vs. 0.0-0.2% sequences), and *Sediminibacterium* (Chinitophagales) was more frequently detected in BV than in other sites (2.5-3.1% vs. 0.0-0.1% sequences; Fig. 7A).

**Figure 7.**
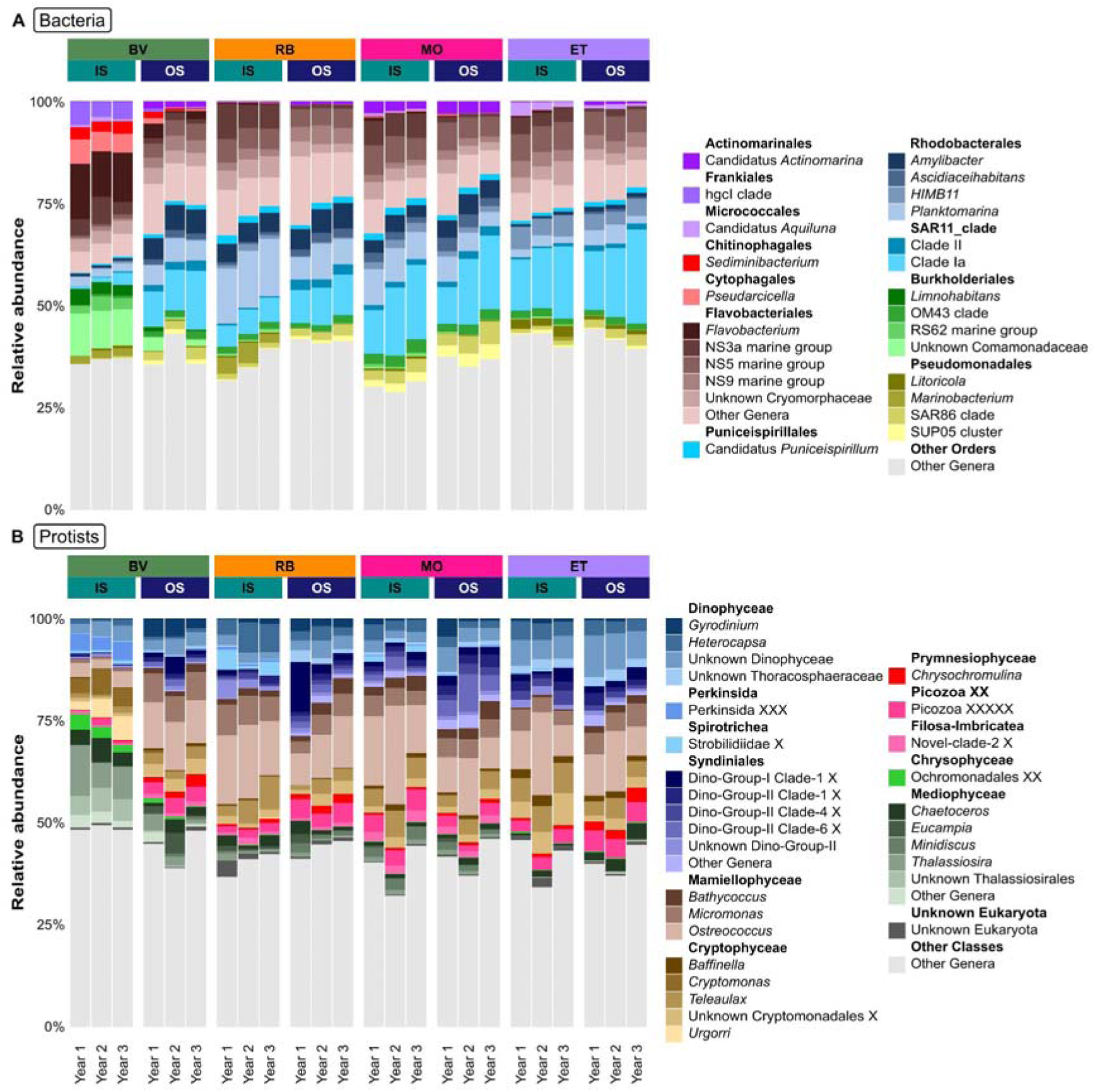
Taxonomic composition of the most abundant bacterial (A) and protist (B) genera in inshore and offshore waters at the four sites for the 3 sampling years. For each coastal site, the 10 most abundant bacteria (A) and protist (B) genera per water type were selected and their relative sequence abundance was represented. The obtained genera were ordered by phylum and order for bacteria (A) and by division and class for protists (B). Less abundant genera were summed and represented as “Other taxa”. Taxonomic groups with a name ending with at least a “_X” were created by PR2 taxonomic experts to homogenize the taxonomic classification. Taxonomic groups identified as “Unknown_” were not taxonomically identified to the corresponding taxonomic level using the PR2 database. IS, inshore waters; OS, offshore waters. BV, Bay of Veys; RB, Bay of Brest; MO, Marennes-Oléron; ET, Thau lagoon. Sampling years were defined as follows: Year 1 corresponds to September 2020–August 2021, Year 2 to September 2021–August 2022, and Year 3 to September 2022–August 2023.

Within the protist community, the most dominant classes were Mamiellophyceae, Dinophyceae, Syndiniales, Cryptophyceae and Mediophyceae accounting for about 3%-31%, 4%-23%, 1.5%-29%, 2%-19% and 1.6%-26% sequences per year, water type and site, respectively (Appendix Table 3B). In OS waters, the most abundant Mamiellophyceae genera among sites and years were *Ostreococcus* (about 7-15% sequences), *Micromonas* (3-7%), and *Bathycoccus* (1-4%, Fig. 6B). Dinophyceae genera were distributed differently in the four sites with *Heterocapsa* presenting higher sequence abundances in ET (about 3% sequences), MO (about 2%) and RB (about 2-3%), and *Gyrodinium* in BV (2-4%), RB (2-3%) and MO in Year 1 (4%). Two Cryptophyceae taxa, *Teleaulax* and Unknown_Cryptomonadales_X, were consistently more abundant in all sites and years accounting for 1-5% and 1-4% sequences, respectively. Patterns of distribution of Syndiniales genera were specific to each site and year with, for instance, higher sequence abundances of Dino-Group-I-Clade-1_X in BV on year 2 (4% vs. 1% the 2 other years) and year 1 at RB (12% vs. 2% the other years). Mediophyceae genera also showed specific patterns with, for instance, *Chaetoceros* being the most abundant genus in ET (2-4% sequences) and showing temporal fluctuations at the other sites. In IS waters, a pattern also observed with bacteria, genera of the five most abundant classes were distributed differently between BV and the three other sites (Fig. 7B). The Mamiellophyceae *Ostreococcus* (7-24% sequences) and *Micromonas* (3-8%) were clearly the most abundant genera in ET, MO, RB while this was only the case of *Ostreococcus* in BV (2-4%). Dinophyceae *Heterocapsa* and Unknown_Dinophyceae were dominant in the three sites (2-7% and 2-6% sequences, respectively) while only Unknown_Dinophyceae in BV (2-4%). Syndiniales genera were mostly observed in ET, MO and BV and presented temporal fluctuations. Cryptophyceae *Teleaulax* and Unknown_Cryptomonadales_X, were most abundant in the three sites (2-8% and 1-8% sequences, respectively) while *Cryptomonas* and *Urgorri* (4-6% and 2-6% sequences, respectively) in BV. The Mediophyceae genera *Chaetoceros*, *Thalassiosira*, Unknown_Thalassiosirales, and *Skeletonema* showed high abundances in BV (6-12%, 3-6%, 5-6%, 2-3% sequences, respectively) but were less noticeable in the three other sites.

Human enteric viruses were investigated in IS (n=12) and OS (n=12) water samples collected from September 2020 to September 2021. As shown on Supp Figure 4 despite purification and DNAse treatment the percentage of viral reads recovered was very low, for both water types (IS: 0.5% and OS: 0.6%). This low percentage was not related to the coastal sites, despite a slightly higher proportion for BV (BV:0.9%; RB 0.3%; MO 0.5% and ET 0.3%). The viral reads obtained did not allow contig assembly and thus no identification of human RNA viruses. The majority of reads obtained from water samples were identified as of bacterial origin, followed by unclassified reads.

Dominant bacteria were different in oysters than in water samples. Those included the orders Enterobacteriales, Mycoplasmatales, Spirochaetales, and Fusobacteriales, representing about 12.9%-38.3%, 10.2%-28.4%, 4.1%-24.1%, and 4.0%-18.4% sequence per year and site, respectively (Appendix Table 3A). Among the most abundant genera in oysters, *Mycoplasma* (Mycoplasmatales), *Vibrio* (Enterobacteriales), and unknown Spirochaetacaeae accounted for 8.4%–22.8%, 3.9%–25.1%, and 3.6%–24.0% sequences, respectively. *Mycoplasma* was more frequently detected at BV than at the other sites (17.3-17.5% vs. 6.9-13.6% sequences; Fig. 8A) while *Vibrio* was more prevalent at ET (17.7-22.3%) and the third year at MO (15.4% sequences) than at the other sites (3.1-10.7%). Finally, sequences affiliated with unknown Spirochaetacaeae were found more prevalent in both RB (8.6-19.6%) and MO sites (9.2-22.9%).

**Figure 8.**
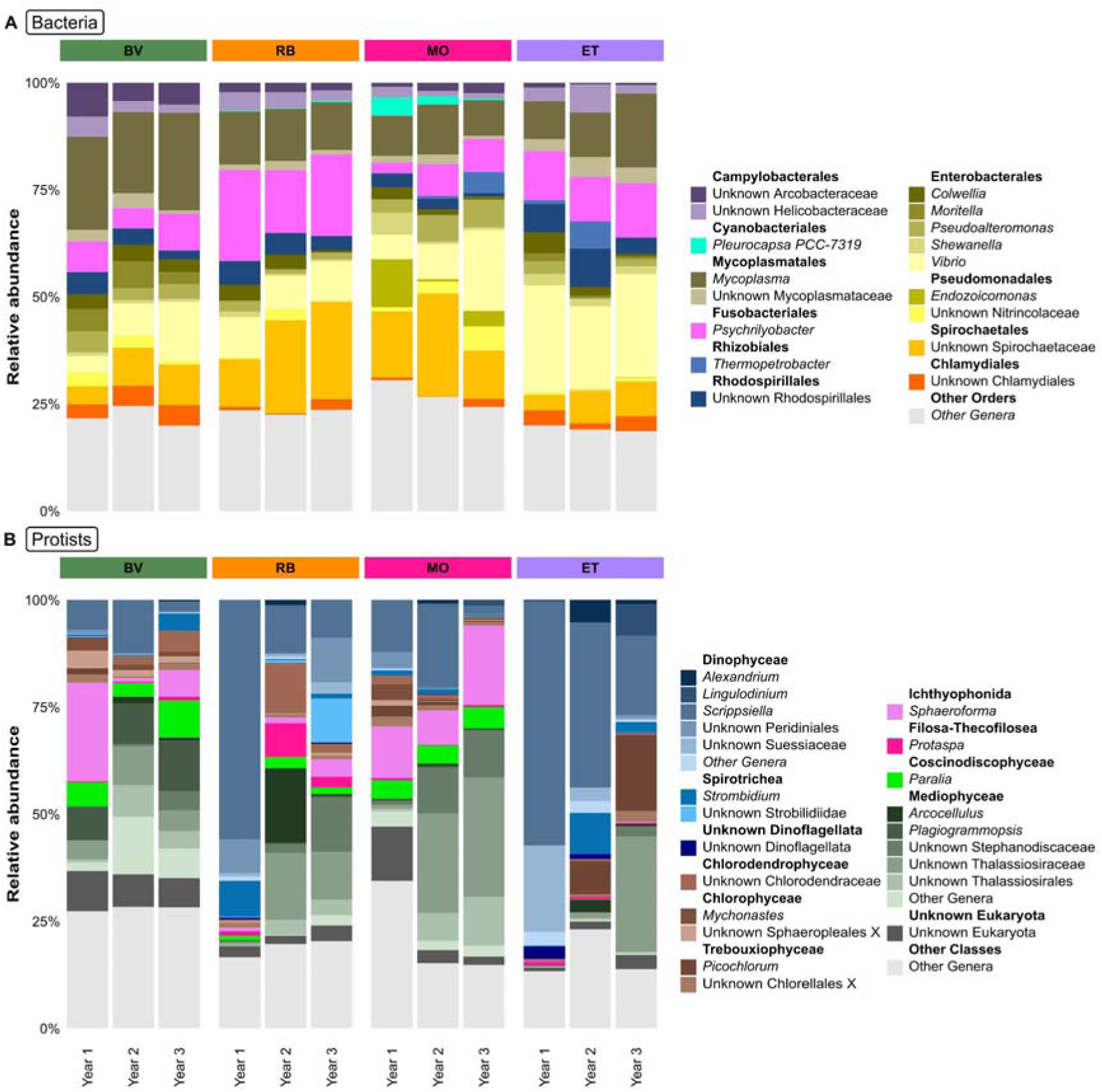
Taxonomic composition of the most abundant bacterial (A) and protist (B) genera in oyster tissues at the four sites for the 3 sampling years. For each coastal site, the 10 most abundant bacteria (A) and protist (B) genera were selected and their relative sequence abundance was represented. The obtained genera were ordered by phylum and order for bacteria and by division and class for protists. Less abundant genera were summed and represented as “Other taxa”. Taxonomic groups with a name ending with at least a “_X” were created by PR2 taxonomic experts to homogenize the taxonomic classification. Taxonomic groups identified as “Unknown” were not taxonomically identified to the corresponding taxonomic level using the PR2 database. BV, Bay of Veys; RB, Bay of Brest; MO, Marennes-Oléron; ET, Thau lagoon. Sampling years were defined as follows: Year 1 corresponds to September 2020–August 2021, Year 2 to September 2021–August 2022, and Year 3 to September 2022–August 2023.

Protist community composition was also different between oyster and water samples. The two most dominant classes were Mediophyceae and Dinophyceae representing about 1%-57% and 4%-86% sequences, respectively (Appendix Table 3B). These classes were followed by Ichthyophonida, Coscinodiscophyceae, Spirotrichea, Unknown_Eukaryota, and Chlorophyceae and that accounted for 1-24%, 2-17%, 2-15%, 2-13%, and 0.4-13% sequences per year and site, respectively (Appendix Table 3B). Dinophyceae reached higher sequence abundances the first year in both RB (69.2%) and ET (85.72%). The most abundant genera in oyster tissues were *Scrippsiella* (Dinophyceae), Unknown_Thalassiosiraceae (Mediophyceae), and *Sphaeroforma* (Ichthyophonida), accounting for about 2–56%, 0.6–28%, and 0.8%–23% sequences across samples, respectively. The genus *Sphaerophorma* (Ichthyophonida) was more abundant in BV the first and third years (23% and 6.3% sequences, respectively) and MO the three years (2–19%). The protist community in oysters from ET was composed of some genera reaching higher sequence abundances such as *Picochlorum* (Trebouxiophyceae, reaching about 18% sequences) and unknown Suessiaceae (Dinophyceae, reaching about 20%) compared to other sites. Furthermore, *Mychonastes* (Chlorophyceae) was detected in all sites but ET. Unknown_Chlorodendraceae (Chlorodendrophyceae) and Unknown_Sphaeropleales_X (Chlorophyceae) did not exceed 0.44% in ET while they reached higher sequence abundances in other sites, such as in BV where these two taxa reached 4.8% and 4.1% sequences, respectively (Fig. 8B).

Human enteric viruses were also investigated in oysters collected from September 2020 to September 2021 by specifically targeting the digestive tissues of the shellfish. As for the water samples, a low proportion of viral reads in the libraries was obtained from these tissue samples (0,3%) regardless of the coastal site. Most of the reads were identified as Eukaryota (61%), followed by unclassified (33%) despite multiple steps to eliminate other sources of nucleic acid than viral RNAs during extraction (i.e. centrifugation, filtration, DNase treatment; Supp Fig. 5). As for the water samples, direct read mapping does not allow the assignment of these sequences to known viral families in the databases used in the MAEVA pipeline. This lack of assignment prevents the estimation of alpha diversity, as well as comparisons of viral communities across sites, years, or water types. Contig assembly of these reads using metaSPAdes also did not provide any reliable or exploitable assignment.

### Pathogen detection

Out of the 314 bacterial genera with members potentially pathogenic to animals, humans, or both, screened in this study, 101 genera were detected in more than one sample from IS waters, OS waters, and/or oyster tissues from the various sites (Appendix Table 2).

Among sequences of genera with bacteria potentially pathogenic to animals, particularly to oysters, sequences of *Cytophaga, Aliiroseovarius* and *Roseovarius* were detected at the four sites. While *Cytophaga* were more particularly detected in water samples, the two last ones appeared more abundant in oysters (Fig. 9A). *Vibrio, Francisella, Mycoplasma*, and *Alteromonas* were repeatedly detected at all sites, in both water and oyster samples. Whereas *Alteromonas* and *Francisella* sequences were detected at low sequence abundances and equally in all samples, *Vibrio* and *Mycoplasma* displayed a much higher abundance in oysters compared to water samples, showing the highest relative abundance in ET oysters for *Vibrio* and in BV oysters for *Mycoplasma*.

**Figure 9.**
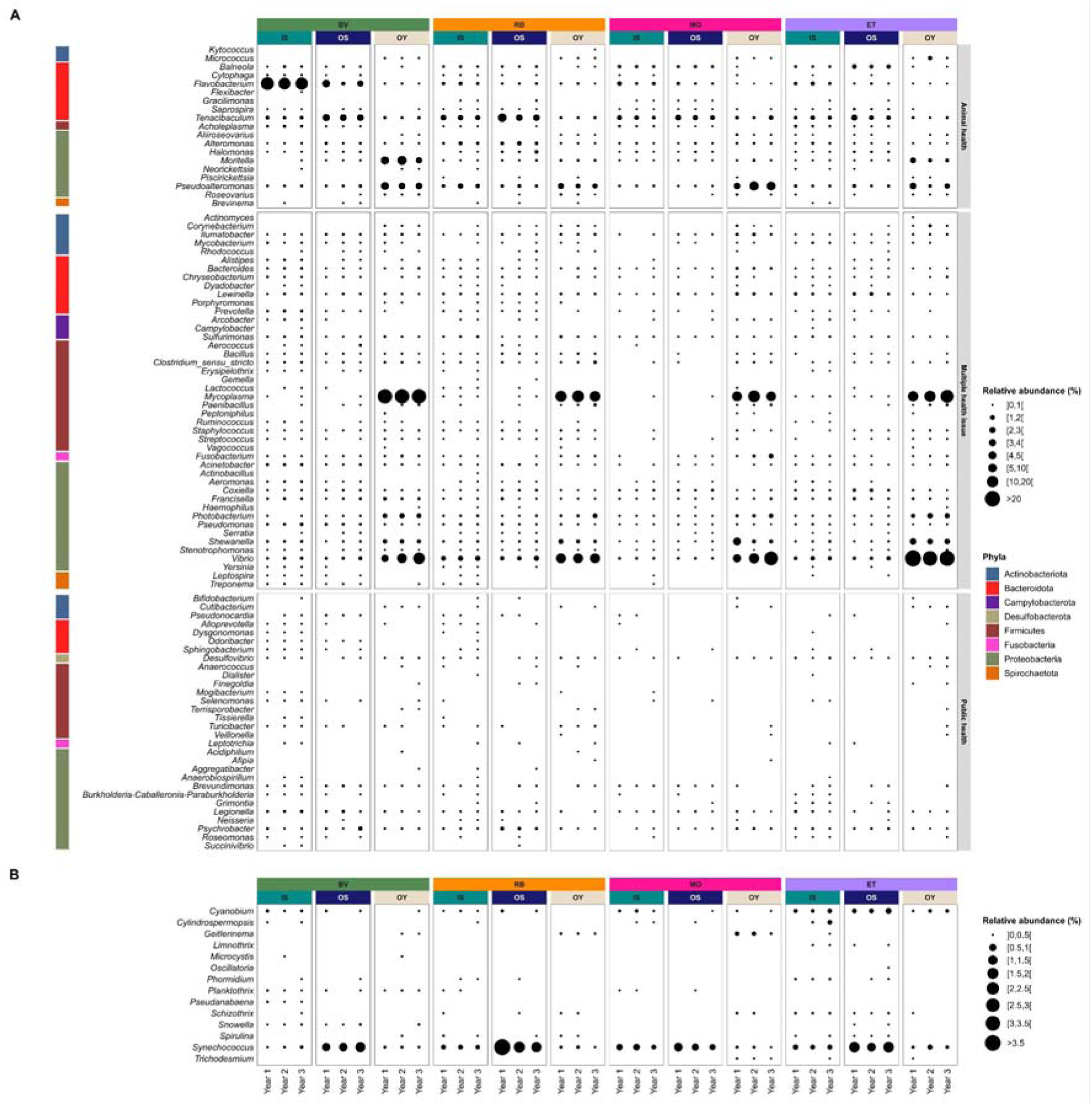
Relative sequence abundance of ASV related to bacterial genera with members potentially pathogenic to animals (animal health), humans (public health), or both (multiple health issues) (A) and to cyanobacterial genera screened for potentially toxic species (B) in the different samples from the four sites for the 3 sampling years.

Together with *Photobacterium*, also a member of the Vibrionaceae family, *Shewanella*, *Vibrio* and *Mycoplasma* represented the most frequently detected marine genera that include human pathogens, particularly in oyster samples (Fig. 9A). Other genera containing bacteria pathogenic to humans but typically associated with freshwater or estuarine environments, including *Legionella, Pseudomonas, Mycobacterium*, and *Clostridium*, were also detected, but at lower frequencies (Fig. 9A). *Legionella, Pseudomonas*, and *Mycobacterium* occurred mostly in water samples, whereas *Clostridium* was mainly detected in oysters. Genera commonly associated with wastewater sources, such as *Acinetobacter*, *Bacteroides, Prevotella, Aeromonas,* and *Arcobacter*, were also detected, particularly in BV-IS samples. *Bacteroides* was the most frequently observed, *Arcobacter* occurred sporadically, and *Aeromonas* was restricted to water samples. In contrast, members of the Enterobacteriaceae family, including faecal bacteria such as *Escherichia coli* and *Salmonella*, were not detected in any IS, OS, or OY samples, in agreement with the absence of human enteric viruses detection.

Among the 67 cyanobacterial genera screened for potentially toxic species, 14 were detected in this dataset. *Cyanobium, Geitlerinema, Limnothrix, Oscillatoria, Schizothrix, Spirulina*, *Synechococcus, Phormidium* and *Trichodesmium*, genera known to occur across a broad salinity range, from freshwater to marine environments, were identified in both IS and OS waters, as well as in OY samples. *Synechococcus* was the most abundant genus overall, particularly in OS waters at all four sites. Freshwater-associated genera, including *Cylindrospermopsis, Microcystis*, *Planktothrix*, *Pseudanabaena*, and *Snowella,* were also detected. Overall, these cyanobacteria were generally found in low or very low sequence abundance in both water samples and oysters (Fig. 9B).

Pathogenic eukaryotic taxa were grouped into different categories according to the type of effects considered. Concerning HAB taxa, 21 genera and 1 family were identified across the entire dataset (Fig. 10). Among taxa associated with harmful effects on environmental health, ASVs assigned to the genera *Heterocapsa* (Dinophyceae), *Chrysochromulina*, *Phaeocystis* and *Prymnesium* (Haptophyceae) were detected in water samples (IS and OS) from all four sites and years, indicating their recurrent presence in the communities (Fig. 10). Within this group, both *Heterocapsa* and *Prymnesium* were less frequently observed in OY samples (Fig. 10). The ichthyotoxic genus *Pseudochattonella* (Dictyochophyceae) was detected infrequently, mainly in OS waters of BV, RB and ET, and in both IS and OS samples at MO (Fig. 10). Regarding Raphidophyceae, *Heterosigma* was present in both IS and OS waters from BV, RB and MO but absent from ET, whereas *Chattonella* occurred exclusively at ET (Fig. 10). The dinoflagellate genus *Margalefidinium* was found predominantly in IS and OS waters at ET, while *Akashiwo* and *Luciella* were detected only in OY samples at BV, MO and ET, and at all sites, respectively (Fig. 10).

**Figure 10.**
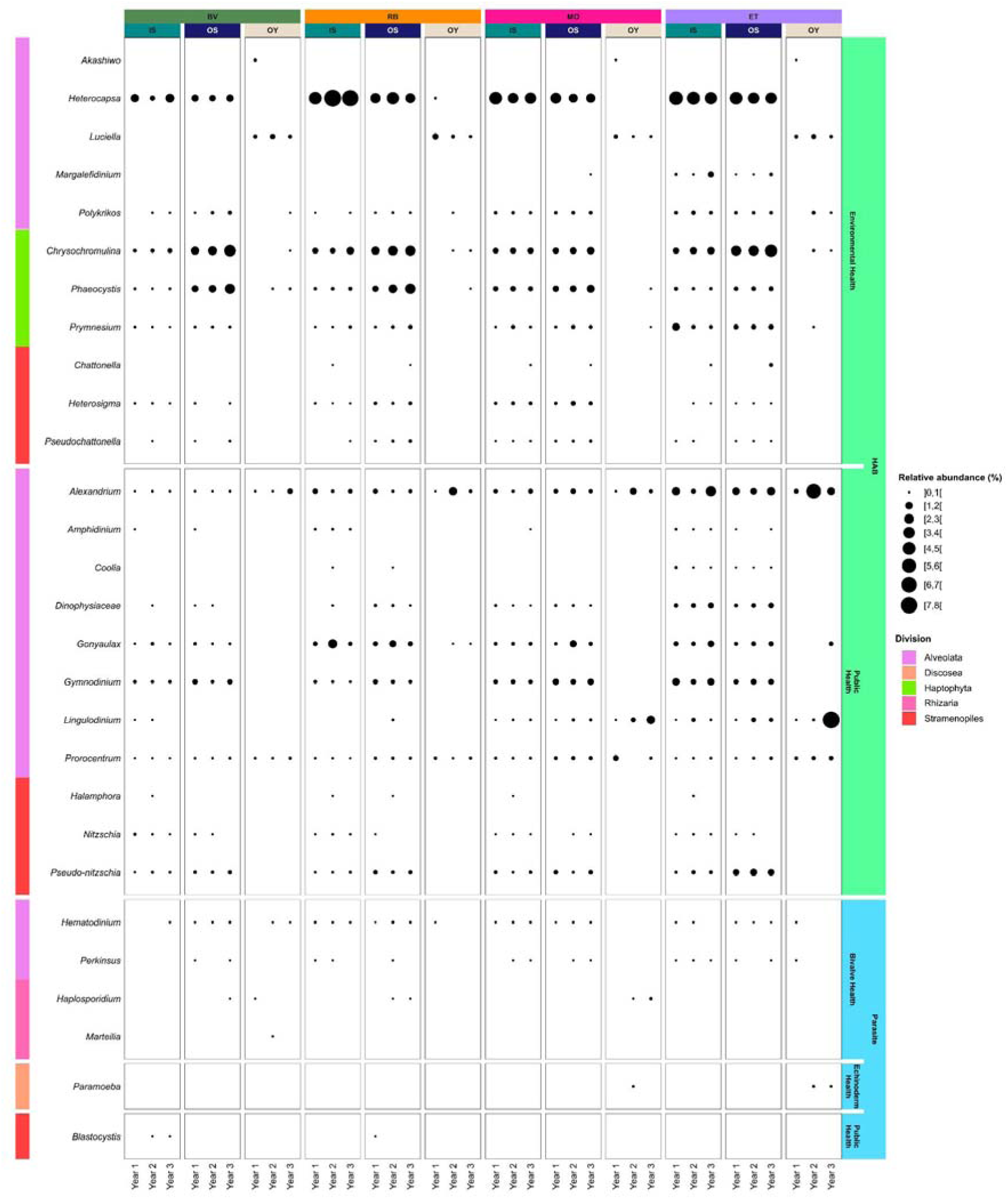
Relative sequence abundance of ASV related to eukaryotic genera (or families) known to include potential HAB or parasites to marine invertebrates or humans in the different samples from the four sites for the 3 sampling years.

Regarding taxa with noxious effects on public health, ASVs assigned to the genera *Alexandrium*, *Gonyaulax*, *Gymnodinium*, *Prorocentrum* (Dinophyceae) and *Pseudo-nitzschia* (Bacillariophyceae) were identified in IS and OS water samples at all sites and years (Fig. 10). While *Alexandrium* and *Prorocentrum* were also detected in OY samples at the different sites each year, other taxa were infrequently observed or not detected at all in OY samples (*Pseudo-nitzschia*). Unexpectedly, the toxigenic genus *Dinophysis* was detected in only one sample, although the family Dinophysiaceae (3 ASVs) appeared at all sites and in all water types. Other taxa, such as the dinoflagellate genera *Amphidinium* and *Coolia*, were more restricted, occurring mainly at RB and ET. Among diatoms, the genus *Nitzschia* was frequently observed at all sites, whereas *Halamphora* was more common in year 2 (Fig. 10).

Among genera known to include bivalve pathogens, no sequence related to *Bonamia* or *Mikrocytos* was detected over the study period in water and oyster samples. Few *Marteilia* sequences were detected in one oyster sample from BV. In contrast, sequences related to *Haplosporidium* and *Perkinsus* genera were detected in several geographic zones and different environmental compartments. While *Perkinsus* was more represented in water than oyster samples whatever their water type, OS or IS, *Haplosporidium* was detected in some OS water samples and oyster samples, particularly from MO. Among the additional genera screened in the sequence data sets, the genus *Hematodinium*, recognized as pathogenic for crustacean, was detected in OS, IS, and some oyster samples. It appeared widely distributed geographically and in time. Interestingly, the genus *Paramoeba*, known to include some sea urchin pathogens, was found in oyster samples from ET and MO. Finally, sequences of *Blastocystis*, a genus including human and terrestrial animal pathogens, were detected in water samples from RB and BV (Fig. 10).

## Discussion

In this paper, we present the first results of the ROME (*Réseau d’Observatoires de Microbiologie Environnementale intégrée*) network which was the first pilot project in France and Europe aimed at developing a national observatory system based on environmental DNA (eDNA) data. Since its conception in 2019, the monitoring system has been designed to integrate the study of different microbiological compartments—bacteria, protists and viruses—by leveraging the potential of nucleic acid analyses. ROME was grounded on the principles of the One Health concept (Wildlife Conservation Society, 2004), collecting data towards the sustainable protection of human health, ecosystems, and biologically exploited resources. The knowledge and data generated form the basis of a 3-year eDNA observatory system that addresses both fundamental research questions and the needs of policymakers and stakeholders. These results are here commented together with the steps made by the ROME project towards the development of a national eDNA-based observatory system.

### Local variability of environmental coastal microbiomes

Our water sampling strategy was to target several planktonic size fractions (i.e. picoplankton and free-living bacteria, 0.2-3 µm; nanoplankton, particle-attached bacteria and larger size fractions, >3 µm; and microplankton and larger size fractions, >20 µm). The aim of targeting the 0.2-3 µm and >3 µm were rather to survey the microbial communities while the >20 µm was to survey microphytoplankton and metazoan groups and their associated microbial communities (i.e. holobionts). Microbiomes data of all size fractions showed separation by size classes, a pattern previously demonstrated in similar studies on coastal ecosystems (de Vargas et al., 2015; Massana et al., 2015; Ramond et al., 2019; Urvoy et al., 2022; Meyneng et al., 2024). For example, members of the SAR11 clade were markedly enriched in the 0.2–3 µm fraction, whereas bacteria from the phylum Bacteroidota (e.g., Flavobacterales, Chitinophagales) and γ-proteobacteria such as Enterobacteriales were more abundant in the > 3 µm fraction. In addition, the diversity analyses confirmed that the results from the >3 µm and >20 µm size fractions were partly redundant. To avoid data redundancy and biased analyses, we decided to remove the >20 µm size fractions from the analyses in this manuscript and keep the 0.2-3 µm and >3µm size fractions (Fig. 2).

Local microbiome variability emerged as a major pattern. When comparing the four sampling sites, both bacterial and protist ASV richness varied (Fig. 3A), and the corresponding cluster of samples were significantly different (Fig. 4). This pattern broadly followed water masses, comparable climatic conditions and geographic latitude. For example, microbiomes from the Bay of Biscay (MO and RB) were more similar to each other than to those from the English Channel (BV) or the Mediterranean Sea (ET). Local microbiome difference across estuarine gradients is commonly observed in coastal ecosystems, with chlorophyll-a concentrations or microscopy-based data (Cottrell et al., 2010; Bilbao et al., 2023), both in association with salinity decreases (Fortunato et al., 2012; Campbell and Kirchman, 2013; Mason et al., 2016; Kirchman et al., 2017; Wu et al., 2024) and classical biogeochemical parameters gradients (Sharp et al., 2009; Crump and Bowen, 2024). Metabarcoding data produced in this study, which broadened taxonomic resolution by providing more diversity data (ASVs), support this pattern and delineate more precise differences among coastal sites.

The difference in microbiomes between inshore vs. offshore water type (IS vs OS) was identified in each studied ecosystem, yet local variability was also evident in this pattern (Fig. 5). In the Bay of Veys (BV), for bacteria and protists, samples from IS clustered apart from OS. These observations are linked to differences in environmental conditions between water types, with OS having a salinity varying between 31.7-34.3 and a temperature between 6.7-21.7 °C, and IS having a salinity varying between 0.2-30.8 and a temperature between 4.1-24 °C over the three years. These differences in diversity patterns between IS and OS were further confirmed by studying their taxonomic composition over the three years, for both 16S and 18S rDNA (Fig. 7). IS samples were dominated by the bacterial genera *Limnohabitans*, *Flavobacterium*, *Pseudarcicella*, and hgcl clade, and by diatoms (i.e. *Chaetoceros*, *Thalassiosira*, Unknown_Thalassiosirales, and *Skeletonema*), Mamiellophyceae (*Ostreococcus*) and Cryptophyceae (i.e. *Cryptomonas* and *Urgorri*) which are commonly identified in freshwaters or waters with low salinities (Marshall et al., 2005; Simon et al., 2015; Mason et al., 2016; Savio et al., 2019; Urvoy et al., 2022; Meyneng et al., 2024). OS waters were mostly dominated by different bacterial taxonomic groups, such as the SAR11 clade, Rhodobacterales (*Amylobacter* and *Planktomarina*), and Flavobacterales (NS5 and NS9 marine groups), and the protist taxa Mamiellophyceae (*Ostreococcus*, *Micromonas*, and *Bathycoccus*), Dinophyceae (*Gyrodinium*), and Cryptophyceae (i.e. *Teleaulax* and Unknown_Cryptomonadales_X). These taxonomic groups are also commonly found in other temperate coastal waters (Gilbert et al., 2010; Massana et al., 2015). The other three sites showed less differences in community structure between the two water types. In Marennes-Oléron and the Thau lagoon, diversity patterns from the two water types were nearly overlapping (Fig. 4, Fig.5). This may be related to the properties of their respective IS and OS water masses which are more similar to each other than in the case of BV (MO: salinity of IS 17-35.5 vs. OS 25.7-35.2 and temperature of IS 6.3-23.6 °C vs. OS 6.8-22.5 °C; ET salinity of IS 33.8-42 vs. OS 35.7-41.2 and temperature of ET : IS 3.9-29.9 °C vs. OS 5.3-28.1 °C). This pattern is shown for instance by the orders SAR11 clade (i.e. clade Ia), Rhodobacterales (i.e. *Amylobacter* and *Planktomarina* for MO, HBM11 and *Planktomarina* for ET), Flavobacterales (i.e. NS5 and NS9 marine groups), Pseudomonadales (i.e. SAR86) that dominated OS and IS bacterial communities. One genus from Flavobacterales (i.e. NS3a marine group) was notably more abundant in IS waters. Regarding the protists, IS and OS waters were both dominated by genera from the orders Mamiellophyceae (i.e. *Ostreococcus* and *Micromonas*), Dinophyceae (i.e. *Heterocapsa*), and Cryptophyceae (i.e. *Teleaulax* and Unknown_Cryptomonadales_X). Still, some differences in genera dominance could be observed between water types: *Bathycoccus* (Mamiellophyceae) dominated OS waters, while Unknown_Dinophyceae dominated IS waters. *Chaetoceros* (Mediophyceae) and Dino-Group-II-Clade-6_X (Syndiniales) dominated OS waters from ET and MO, respectively. Our observations are in line with previous findings made by eDNA analyses, such as a 6-months survey of monthly-taken water samples targeting bacteria and protists in the 0.2-3 µm fraction in the Bay of Biscay (Garate et al., 2022). In this last study, samples from a coastal site and an offshore site indicated that although the most abundant taxa followed similar temporal trends, some taxa were specifically more abundant in each of the sampling sites.

In general, the origin of community differences between inshore vs. offshore sites in water eDNA studies has been linked to environmental conditions (e.g., water temperature; Hervé et al., 2025; salinity; Herlemann et al., 2011; Urvoy et al., 2022), including continental pressures that influence coastal ecosystem biodiversity (Zoppini et al., 2019). Among these, river runoff is a significant factor associated with shifts in estuarine biodiversity (Fagervold et al., 2014; Urvoy et al., 2022; Meyneng et al., 2024). River discharge can introduce freshwater and terrestrial species and cause salinity drops, which facilitate the development of new species while leading to the decline of others (Jiajun et al., 2024; Amadei Martínez et al., 2025). Our study directly assessed the impact of river runoff on coastal microbiome diversity, using a comparative approach between stations subject to varying degrees of riverine influence. Although large French estuaries such as the Gironde, Loire, and Rhône were not included in the present study, it can be supposed that stronger river flow likely accentuates the coastal-offshore biodiversity gradient as already observed in other systems such as the Mississippi estuary (Mason et al., 2016), the Seine estuary (Hervé et al., 2025) or the Pearl River Estuary (Wang et al., 2021).

Interestingly, coastal-offshore differences were observed both in tidal bay (RB) and non-tidal lagoon (ET), a more enclosed ecosystem and with less river input. We demonstrated that the coastal-offshore biodiversity difference is closely linked to river flow intensity and the location of the sampling station along the river continuum. Nonetheless, the spatial positioning and the number of sampling stations remains critical to properly characterize this gradient. The further upstream the sampling location, the greater the variability we observed in ASV composition (e.g., BV, Fig. 5). In contrast, when river flow was weaker and tidal dynamics created reverse estuary conditions (e.g., MO, Fig. 5) (Latouche and Jouanneau, 1990) - where marine waters enter river mouths - differences in coastal-offshore biodiversity were minimal.

While local microbiome biodiversity and coastal-offshore biodiversity gradient has been described in previous studies, these were often limited in duration—sometimes to a single sampling campaign and/or only one sampling site (Fortunato et al., 2012; Tee et al., 2021; Wang et al., 2021; Hervé et al., 2025). Our study combines the advantages of high-resolution metabarcoding data at two sampling stations with contrasting water types with longitudinal sampling across three years and three distinct estuaries and one lagoon. This showed that local microbiome variability in estuaries and the coastal-offshore microbial community structuration is a persistent, though dynamic, feature of estuarine ecosystems.

### Oyster and planktonic microbiomes intercomparison

One major innovation of the ROME project was the collection of organismal microbiome eDNA from oysters in parallel with environmental planktonic microbiomes. In alignment with the One Health frame. This strategy was designed to analyze changes in the oyster microbiome in relation to local ecosystem variability, based on the idea that environmental conditions are key drivers of microbiome shifts and that oyster microbiome variability could indicate environmental changes (King et al., 2019; Walker et al., 2025).

At the moment, a perfect comparison between oyster-associated microbiomes and environmental microbiomes by metabarcoding approaches is challenging due to the lack of common methods to amplify microbial communities from water and oyster samples. In fact, the primers targeting protists in oysters (Clerissi et al., 2018) were specifically designed to avoid the amplification of host DNA, which could otherwise limit a comprehensive assessment of oyster-associated microbiomes. Similarly, a primer set targeting bacteria in oysters (Klindworth et al., 2013) was selected to preferentially amplify bacterial DNA and to prevent amplification of some eukaryotes (e.g., protists) present in oysters, which could be amplified by the primers of (Parada et al., 2016), as observed by (Walker et al., 2025) in whole oysters. Furthermore, oyster samples were collected monthly, with a slight delay relative to the biweekly water sampling, introducing a temporal mismatch. Despite these limitations, in our study, we suggest that oysters can be valuable ecological sentinels for monitoring environmental variability since they can show local microbial variability of the sampling sites, as previously observed (Walker et al., 2025). Our study clearly demonstrated that oyster microbiomes vary locally as environmental microbiomes and it is known that oyster microbiome is highly influenced by both environmental and host-related factors (Paillard et al., 2022). In our study, the observed site-specificity of the oyster microbiome communities suggests the presence of selective processes in the local oyster populations in addition to local environmental factors that resulted in local variability of microbial oyster assemblages. This result was achieved thanks to the multi-year and multi-site datasets of oyster-associated microbiomes in conjunction with environmental variability of the project ROME. Large scale and multiyear datasets are rare, and most existing studies focus primarily on human or oyster disease identification rather than ecological patterns (King et al., 2019). This makes the ROME dataset a unique and precious pioneering study to associate oyster and planktonic environmental microbiome variations in relation to environmental variability.

### Detection of potential pathogens

Through the analysis of coastal microbial communities, the ROME project also aimed at evaluating to which extent metabarcoding and metagenomic approaches from a unique sample could be used to detect emerging and previously undetected pathogens. These include human enteric viruses, animal or human pathogenic bacteria, harmful microalgae, and oyster pathogens, which are organisms that often fall outside the detection range of conventional identification methods such as microscopy (for harmful algae) or targeted molecular approaches like qPCR (for oyster pathogens, viruses and bacteria, after a strain culturing).

As expected, the chosen genomics approaches have allowed detecting a large spectrum of genera that include potential human and animal pathogens rather than pathogen species. Nevertheless, no reads of human enteric viruses or bacteria belonging to Enterobacteriaceae, which include the genera *Escherichia* and *Salmonella,* were detected in coastal waters or oysters throughout the monitoring period. The analysis of oyster batches collected from shellfish harvesting areas classified as category B (BV, RB, and ET sites), together with water samples collected near riverine inputs potentially impacted by faecal contamination could have been in favour of the detection of human enteric viruses and enteric bacteria in our samples. For example, in a previous study at other shellfish harvesting areas classified as category B in France, pathogenic enteric bacteria (i.e. *Campylobacter*, *Salmonella*, pathogenic forms of *Escherichia coli*) were isolated and human enteric viruses such as norovirus were detected by PCR in oysters (Rincé et al., 2018).In fact, shellfish samples including oysters are known to be able to accumulate human enteric viruses (Bonny et al., 2021; Desdouits et al., 2021; Schaeffer et al., 2023) and an European study showed that the probability to detect human norovirus is higher in oyster samples collected in contaminated areas (like class B or C) compared to non-contaminated area (EFSA, 2019).

The absence of such pathogens in our samples is likely explainable by the dilution factor of contaminated coastal waters. This hypothesis is supported by previous results showing the rapid dilution of the contamination coming from human sewage which is the main source of virus introduction in estuarine waters (Farkas et al., 2018) and the very low abundance of human enteric bacterial pathogens compared to pathogens from marine origin in mussels located at 175 m from the outfall of a treatment plant in Spain (Ríos-Castro et al., 2023). Furthermore, viruses do not multiply outside their hosts and to our knowledge, have never been evidenced by metagenomics without testing large volumes of water (> 100 L instead of 1 L used in this study). Yet, the detection of some noroviruses using PCR tools in coastal seawater samples using the same concentration methods remains possible (Desdouits et al., 2021). The absence of detection of contigs of human enteric viruses, which are mainly single strand RNA viruses in our oyster samples could be due to the difficulties to detect such small genomes (from 7 to 8 kb) among genomes of other microorganisms present in environmental samples without any enrichment step (Strubbia et al., 2019; Schaeffer et al., 2023; Izquierdo-Lara et al., 2025; Li et al., 2025). Thus further development of the research of human enteric viruses by metagenomics in oysters should include this technical step. Finally, viral databases are limited, resulting in a high percentage of unclassified reads in environmental samples (Nieuwenhuijse et al., 2020). Therefore, we cannot exclude that our negative detection is also due to the limitation of the current databases.

In contrast to Enterobacteriaceae, other bacterial genera known to include human enteric pathogen species such as *Acinetobacter*, *Bacteroides*, *Prevotella*, *Aeromonas*, and *Arcobacter* were detected in water and oyster samples, suggesting episodic contamination by anthropogenic sources (e.g. either from wastewater discharges or from agricultural runoffs). Comparable bacterial detection, except the absence of Enterobacteriaceae, was observed in waters from a shellfish growing area downstream urban and agricultural areas in the United States, where a link with heavy rain was also suggested (Leight et al., 2018). Moreover, marine bacteria including human pathogenic members, such as *Vibrio, Shewanella*, and *Mycoplasma*, were predominantly identified in oysters. The genus *Vibrio* includes species such as *V. parahaemolyticus*, *V. vulnificus*, and *V. cholerae*, known to cause shellfish-borne illnesses (EFSA, 2024).

Concerning potential animal pathogens, many bacterial genera including species pathogenic for fish (e.g. *Flavobacterium, Micrococcus, Pseudoaltermonas, Tenacibaculum*) and also for bivalves (e.g. *Francisella*) were detected in water and oyster samples. Giving that oysters might integrate biodiversity changes over time, these bivalves could be potentially used as sentinels for the surveillance of these pathogens but that would need to be further demonstrated. Considering their potential impact on animal and human health, the detection of *Aliiroseovarius, Roseovarius Cytophaga and Francisella* sequences would need further characterization to understand their presence in our oysters. For example, *Roseovarius crassostreae,* related to *Aliiroseovarius* and *Roseovarius* genera, and some Cytophaga-like bacteria were previously reported in the U.S.A. in the context of oyster mortality and associated with the hinge oyster disease, respectively (Dungan and Elston, 1988; Gómez-Chiarri et al., 2025) Additionally, the detection of *Vibrio* and *Mycoplasma* in high abundance in oysters in all years and sites would deserve complementary investigations to test the potential presence of health indicators or pathogenic taxa as well as their seasonal and spatial dynamics. Indeed, *Mycoplasma* was identified as a potential health indicator taxa along with some *Vibrio* ASVs (Wegner et al., 2025).

Overall, only four genera known to include marine bivalve parasites were identified in our dataset: *Haplosporidium*, *Perkinsus*, *Minchinia*, and *Marteilia*. Very few samples contained sequences of bivalve parasites (fewer than 10 per water type and site). *Haplosporidium* and *Perkinsus* were detected in both oyster and water samples across three to four sites, whereas *Minchinia* and *Marteilia* were only found in oyster samples from a single site. Species belonging to these four genera are known to occur in France, and some, such as *Haplosporidium costale*, are also known to infect the Pacific cupped oyster (Arzul et al., 2022). However, our results do not allow us to determine whether their detection in oyster tissue reflects a true infection or simply passive carriage. In contrast, no sequences of *Bonamia* or *Mikrocytos* were detected in either oyster or water samples, despite reports of these genera in various French sites. This absence may be due to several factors: poor amplification of these species with the selected primers, limitations in the sampling strategy, or biological traits of the parasites that result in a stronger and more consistent association with the host throughout their life cycle (Mérou et al., 2023). Conversely, we detected sequences of *Hematodinium* and *Parameoba*—two genera known to include pathogens of crustaceans and echinoderms, respectively (Buchwald et al., 2018; Alimin et al., 2024). in some oyster tissue samples. While *Parameoba* was found only in oysters, *Hematodinium* was also detected in OS and IS samples across several sites, suggesting widespread distribution. Conversely, *Blastocystis*, a genus previously reported in mussels (Ryckman et al., 2024) was only detected in a few water samples in our study. These results support the interest of using oysters to act as indicators for the presence of species pathogenic to other marine invertebrates or even humans.

Although detected in low abundance, several freshwater cyanobacteria genera were identified. Some of these genera (*Microcystis*, *Planktothrix*, *Pseudanabaena*) have already been reported as potentially toxic taxa along a land-sea continuum in French brittany (Bormans et al., 2019, 2020; Reignier et al., 2024), demonstrating their likely introduction through riverine inputs. Similar transfers have been documented worldwide, most often involving *Microcystis*, the most widespread freshwater microcystin-producing cyanobacterial genus (Preece et al., 2017; Yousaf et al., 2025). Nevertheless, the occurrence of cyanotoxins beyond microcystins (e.g., anatoxins, cylindrospermopsins, nodularins, saxitoxins) in brackish and marine waters, as well as in marine organisms (Biré et al., 2020; Amzil et al., 2023) without a clearly identified source, suggests the presence of additional toxic cyanobacteria (Fraga et al., 2025). Our findings extend these observations by revealing the presence of such potentially toxic taxa in coastal ecosystems. Interestingly, our analysis also revealed the presence of *Trichodesmium*, a dominant diazotrophic genus typically associated with tropical and subtropical regions, in ET and MO samples.

Regarding harmful microalgae, 21 out of 40 genera and 1 out of 3 families of HABs were detected in our study. Among the genera detected, some taxa poorly detected and identified by traditional microscopy methods were present in different sites/water-types such as *Pseudochattonella* (Dictyochophyceae), *Heterosigma* (Raphidophyceae), *Chrysochromulina* and *Prymnesium* (Haptophyceae). On the one hand, while some toxic genera such as *Alexandrium* and *Pseudo-nitzschia* are well retrieved in the dataset, the genus *Dinophysis* was more problematic and ASVs were mostly assigned at the Family level (Dinophysaceae) and almost not at the genus level (1 hit on the whole dataset). On the other hand, some harmful algae belonged to very diversified genera for which only one or a few species are listed as toxic (as for instance in the genera *Gymnodinium, Heterocapsa* or *Nitzschia*). In such cases, positive hits at the genus level do not necessarily imply that a harmful/toxic species is present. The absence of ASVs assigned to the common dinoflagellates in the Kareniaceae (*Karenia* spp. and *Karlodinium* spp.) and Amphidomataceae (*Azadinium* spp.) well known in Atlantic waters (Nézan et al., 2014; Luo et al., 2017) is rather surprising and it may be the result of the sequence assignment method used and poor taxonomic information in the V4 domain for these groups. Regarding HABs sequences in OY samples, only a few of the taxa found in water samples were detected. This is the case of the genus *Luciella* found exclusively in OY samples and not in water samples. An effect of the use of different primers for waters and oysters cannot be excluded to explain such discrepancies, as the primers may have different affinities with targeted taxa. Overall, these results highlight the potential of metabarcoding to complement traditional methods for surveying HAB species diversity, especially highlighting the presence of taxa that were not expected to occur in the studied environment and in French metropolitan areas. When moving to monitoring strategy and regulation application for shellfish, short-read metabarcoding might be limiting for risk assessment for shellfish consumption. A species-specific assignment is absolutely required and qPCR/dPCR approaches might complete a metabarcoding survey in monitoring risks associated with HABs.

### Towards the development of an integrated eDNA-based observatory system

ROME emerged from a balance between scientific ambition and the constraints of internal funding, administrative limitations, and technical constraints. About forty permanent staff members—including technicians, engineers, and researchers—contributed to the project, and compromises were necessary to identify the minimal, yet essential, analyses needed to achieve meaningful results. These decisions enabled the development of a strategy which prioritized bacterial and protist metabarcoding over metagenomics, while metagenomics for human viruses was deemed essential. Sampling efforts (fortnightly for water, monthly for oysters) and sequencing strategies (primer selection, sequencing depth, etc.) were constrained by both budget and human resources and actual technical advancements on the chosen methods at the time. Our approach relied on the scientific division of tasks among national laboratories, which were responsible for sampling, nucleic acid extraction, amplification, metadata management, statistical analysis, and data sharing. We envisioned a flexible and evolving network, supported by the biobanking of original samples (including extracted DNA, half-filters and oyster tissues) to allow for reanalysis as extraction and sequencing technologies advance. Access to samples is governed by shared rules, and all ROME data are freely available to the scientific community beyond institutional boundaries. The ROME project was the first pilot study in Europe aimed at developing an integrative eDNA-based microbial monitoring program for coastal ecosystems. Building on this foundation—and considering the growing need for long-term ecological observations (Cocciardi et al., 2024) —future developments could be envisioned in parallel with ongoing advancements in environmental genomics research and technologies.

Automated sampling and high-throughput sequencing technologies are expected to accelerate workflows and enhance data quality. While eDNA water samplers are under development (e.g.Verdier et al., 2022) these devices are still largely tailored to project-specific needs, and no standardized instrumentation has yet been adopted. Nevertheless, automated sampling could significantly reduce filtering time and human resource requirements, which often constitute problematic issues for long-term monitoring programs. Meanwhile, portable and rapid sequencing technologies such as the Oxford Nanopore Technology (e.g. MinION sequencer), or long-read platforms like PacBio, are becoming increasingly accessible. Ongoing studies comparing short-read (ca. 400 bp) and long-read (ca. 1500 bp) sequencing approaches suggest that longer reads can improve taxonomic resolution. However, reference databases are still predominantly built from short-read barcode sequences targeting hypervariable regions, limiting the potential of long-read data. Thus, improving and continuously updating taxonomic databases remains a major bottleneck for eDNA-based approaches. This calls for coordinated international efforts—bringing together taxonomists, biologists, ecologists, and bioinformaticians—to develop consensus-based, ecologically relevant databases, particularly in regions of the world where marine biodiversity remains poorly characterized.

In ROME, different primer sets were used to target bacteria and protist communities in both water and oyster samples. One of the next logical steps toward optimizing and standardizing eDNA-based monitoring is the validation of universal primers. Some candidate primer sets have been proposed (Priest et al., 2025), but their application at the scale of long-term, multi-site studies raises concerns regarding sequencing depth and associated costs. Additionally, when analysing holobionts—such as oysters—these universal primers may require the use of blocking primers to prevent host DNA amplification. However, such blocking strategies may introduce biases in microbial diversity assessments, and their potential impact needs to be thoroughly evaluated before universal primers can be broadly adopted.

The combination of metabarcoding (for bacteria and protists) and metagenomics (for human viruses) in the ROME project proved useful for assessing community-level biodiversity. However, our approach (i.e. selecting short hypervariable regions of 16S and 18S rDNA) showed limitations in detecting specific pathogens at the species level. These limitations were partly due to the use of short-read sequencing, which restricts taxonomic resolution to the genus level, the selected gene fragment, and to the choice of primers that failed to amplify species of some pathogenic groups (e.g., *Vibrio* spp., *Alexandrium* spp.). In addition, abundance data of pathogens species are essential to assess risks associated with those taxa. Pending future development of the metabarcoding approach towards quantitative estimations, this approach at the moment allows semi-quantitative data in terms of reads of a marker gene which can have different copy numbers depending on microorganism species, as well as several technical limitations at each metabarcoding step. Consequently, targeted quantitative methods such as qPCR or dPCR remain essential for monitoring known pathogenic microorganisms within an eDNA observatory framework. Some pathogens that were expected to be detected in our samples were absent (e.g., *Dinophysis* spp.), which may be due to the specific site selection as well as the targeted gene fragment, likely not specific enough. For example, concentrations of enteric bacteria or human viruses in estuarine water may have been too low to be detected with metabarcoding. Sampling further upstream in the watershed may have revealed different pathogen profiles. Nonetheless, ROME was designed to evaluate pathogen-related risks specifically in estuarine and coastal waters, and our sampling strategy was aligned with this objective. This highlights the importance of carefully designing the spatial and temporal sampling strategy of any eDNA monitoring network to ensure that it meets its intended goals. ROME did not address certain key components of coastal biodiversity, such as the water virome. Only human viruses extracted from oyster tissues and water samples were analyzed. Likewise, important habitats like surface sediments were not included. Yet both the water virome and superficial sediments can host specific microorganisms—including pathogens—that could provide insights into site-specific variability and emerging threats to human health and ecosystem sustainability. Importantly, ROME’s water sampling sites (IS sites) were integrated into the TREC (Traversing European Coastlines) initiative (https://www.embl.org/about/info/trec/), during which soils, sediments, aerosols, surface waters, and offshore marine waters were systematically sampled across 115 sites in Europe. The results from this large-scale exploration will undoubtedly complement and enrich the ROME dataset and its potential.

## Conclusions

At institutional, national, and international levels, ROME marks a step forward toward the standardization of eDNA monitoring, highlighting both the importance and the complexity of establishing such practices for future applications. It paved the way for the development of genomic-based observatory in France and Europe, with the dual aim of advancing research on coastal microbiome dynamics and supporting policy objectives related to the sustainability of exploited resources and the protection of human health. The benefits of this pilot project are evidenced by the results obtained in this study which also addresses potential methodological and strategical improvements in the future.

ROME methods and data production focused on the microbiological compartment, but given the broad applicability of eDNA to the entire biological spectrum, future observatories should expand beyond the microbial domain. These observatory systems should include other marine taxa (e.g., mesozooplankton, fish, marine mammals), whose biodiversity assessment and conservation are central to both scientific inquiry and ecosystem management (e.g., in the context of Marine Protected Areas). This work establishes the foundation for the first analyses of the ROME database, which holds strong potential for further exploitation. Additional studies and data sets exploitation will follow in the future, each addressing specific scientific questions and topics

Here we showed that despite a common structure in terms of size classes separation, alpha diversity and community structure, local ecosystem variability is a defining feature of both water and holobiont-associated (oyster) microbiomes. This suggests that the assessment of the coastal microbiome ecology needs mesoscale analyses to contextualize this variability within the larger framework of coastal ecosystem dynamics. It shows the need for an intercomparable, structured network of coastal observatories to look for common general ecological patterns that overpass local variability.

River runoff was shown to influence coastal microbiome diversity. Bacterial and protist communities varied from inshore to offshore, with the magnitude of these changes depending on river discharge as well as geographic and environmental characteristics. It remains to be investigated whether interactions between microbial compartments are also shaped by this influence, and whether river runoff promotes specific bacteria–protist associations. Based on these gradients, bioindicators of riverine input could be developed and tested across different estuarine systems. Furthermore, the temporal impact of river runoff could be correlated with pressure indicators, such as contaminant levels in water or sediments, to assess whether bioindicators of runoff might also serve as proxies for broader anthropogenic pressures (e.g., industrial or agricultural pollution).

Despite some limitations in detecting specific pathogens, the metabarcoding approach used in ROME proved effective for identifying pathogens unrecognizable by classical monitoring systems based on microscopy. This shows the potential of metabarcoding as a screening biodiversity tool to detect the presence of unsuspected microorganisms in a studied ecosystem. The ROME system thus provides a foundation for the development of an early warning system for environmental and health risk assessment which would be of high relevance for both local and national stakeholders involved in coastal management. The full integration of eDNA data into coastal observatories will require alignment with conventional datasets (e.g. microscopy) and the integration of other eDNA approaches such as the qPCR or dPCR targeted detection. In a future integrative eDNA observatory, a combination of techniques and strategies appears promising: short-read sequencing for community-level ecological monitoring and bioindicator development, alongside long-read sequencing to detect rare or unexpected species in a non-targeted manner, with the addition of targeted approaches (qPCR, dPCR) to quantify species under legislative regulation—such as toxin-producing microalgae, enteric bacteria, and bivalve pathogens—that require precise abundance thresholds to inform mitigation and risk-reduction strategies (e.g., shellfish safety for consumption, bathing water quality, etc.).

## Acknowledgements

ROME was entirely funded by Ifremer in its conception, human resources engagement and costs. The ROME consortium wishes particularly to thank all personnel involved in sampling activity of the coastal laboratory (LER/N, LER/BO, LER/PC, LER/OC) for their constant devotion throughout the duration of the project. A special thanks to Brendan Hennebaut for his help in data and sample biobanking and to Michel Ropert for producing the figure of the map of the sampling sites. Undergraduate students who participated in the ROME project are acknowledged as well as all Ifremer volunteers and colleagues who participated in its logistic and scientific development. ROME is part of the project FUTURE-OBS (ANR-22-POCE-004) of the French Programme Prioritaire (PPR) “Océan et climat”. The database was additionally used in the frame of the co-Funded project Biocean5D (https://biocean5d.org/) and Obama-Next (https://obama-next.eu/). Views and opinions expressed are however those of the author(s) only and do not necessarily reflect those of the European Union or. Neither the European Union nor the granting authority can be held responsible for them.

## Author contribution

RS created, managed and scientifically directed the project ROME since its beginning in 2019. JCP co-managed the project since 2022. RS, IA, NC, AG, CN, MG analysed data and contributed equally in manuscript writing. EB, MC, GC, PD, SLG, SP, CM contributed to data production, analyses and manuscript corrections. THF, JFP, OS, contributed to data acquisition, manuscript writing, field and laboratory expertise. JS contributed to data acquisition, sample biobanking and technical supervision of laboratory procedures. AC, CF, SF, CG, LLB, CL, CM, JQ, SS, JLS, ATT contributed to sampling and/or data acquisition. LL curated data banking and publication.

## Data availability statement

The data that support the findings of this study are openly available in ENA at https://www.ebi.ac.uk/ena, with the project reference number PRJEB105940 and are accessible at https://doi.org/10.12770/a14dedb4-65e8-48dc-a07b-be48d546a5df.

## Ethical statement

The authors declare that the research meets the ethical guidelines of the study country.

## Conflict of interest statement

We declare that the work described has not been published previously, that it is not under consideration for publication elsewhere, that its publication is approved by all authors and by the IFREMER where the work was carried out, and that, if accepted, it will not be published elsewhere in the same form. We disclose any financial, non-financial, and competing interests with other people or organizations that could influence the results and discussion reported in this paper.

## Supplementary figures

**Supplementary Figure 1.**
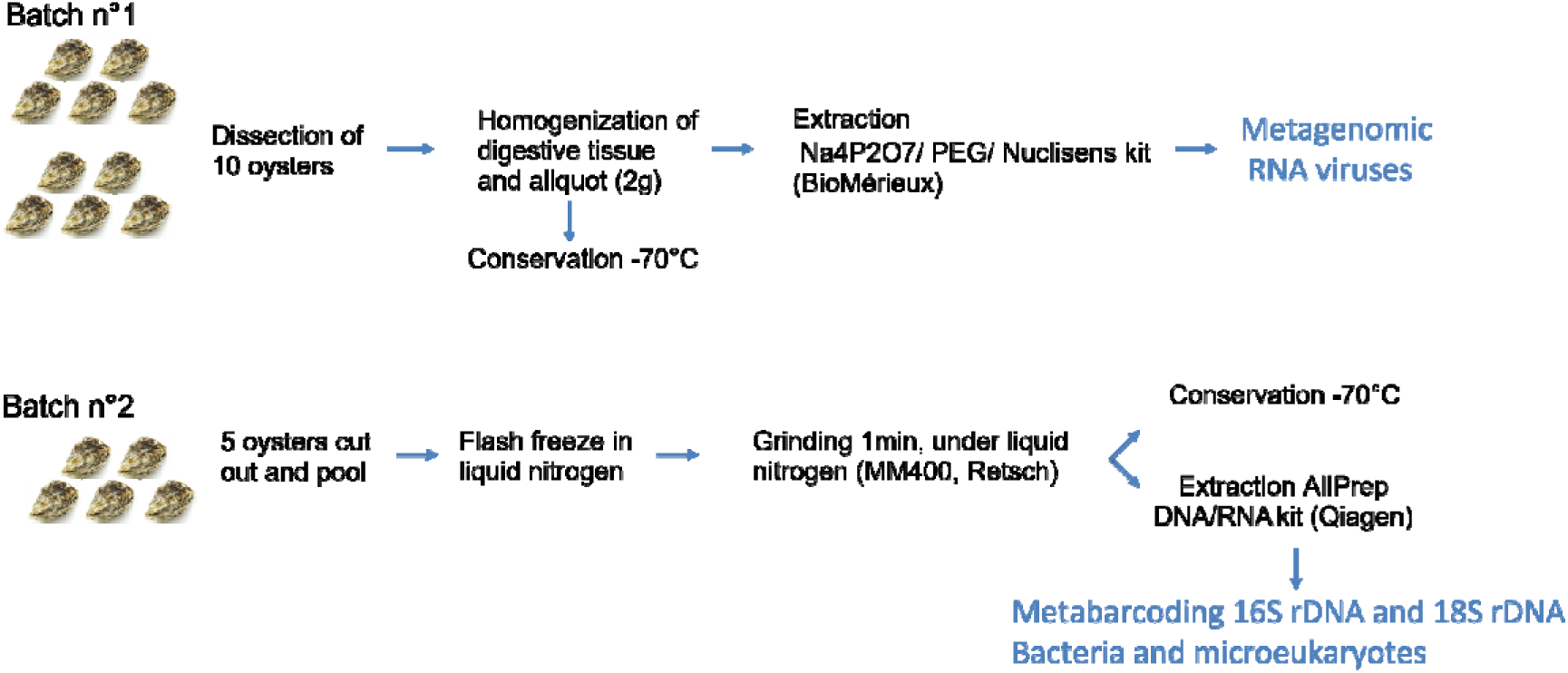
Experimental strategy for oyster analysis, including dissection of digestive tissue (Batch n°1) or retrieval of whole organisms (Batch n°2), sample conditioning (adapted to each batch), DNA extraction and further processing (either metagenomics or metabarcoding).

**Supplementary Figure 2.**
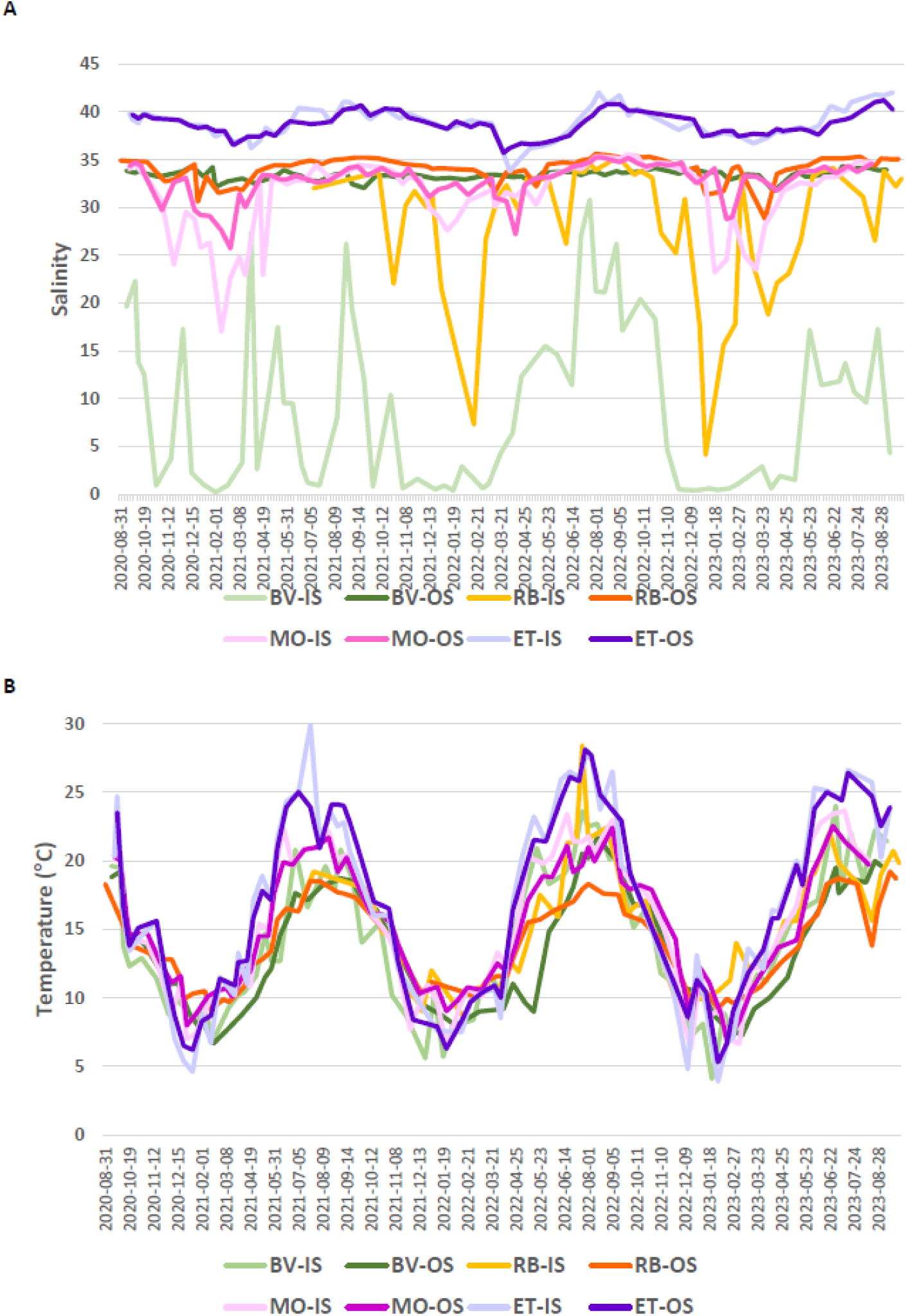
Salinity (A) and temperature (B) measurements in water samples for each site and water type from September 2020 to August 2023. Supp Fig. 2B Temperature

**Supplementary Figure 3.**
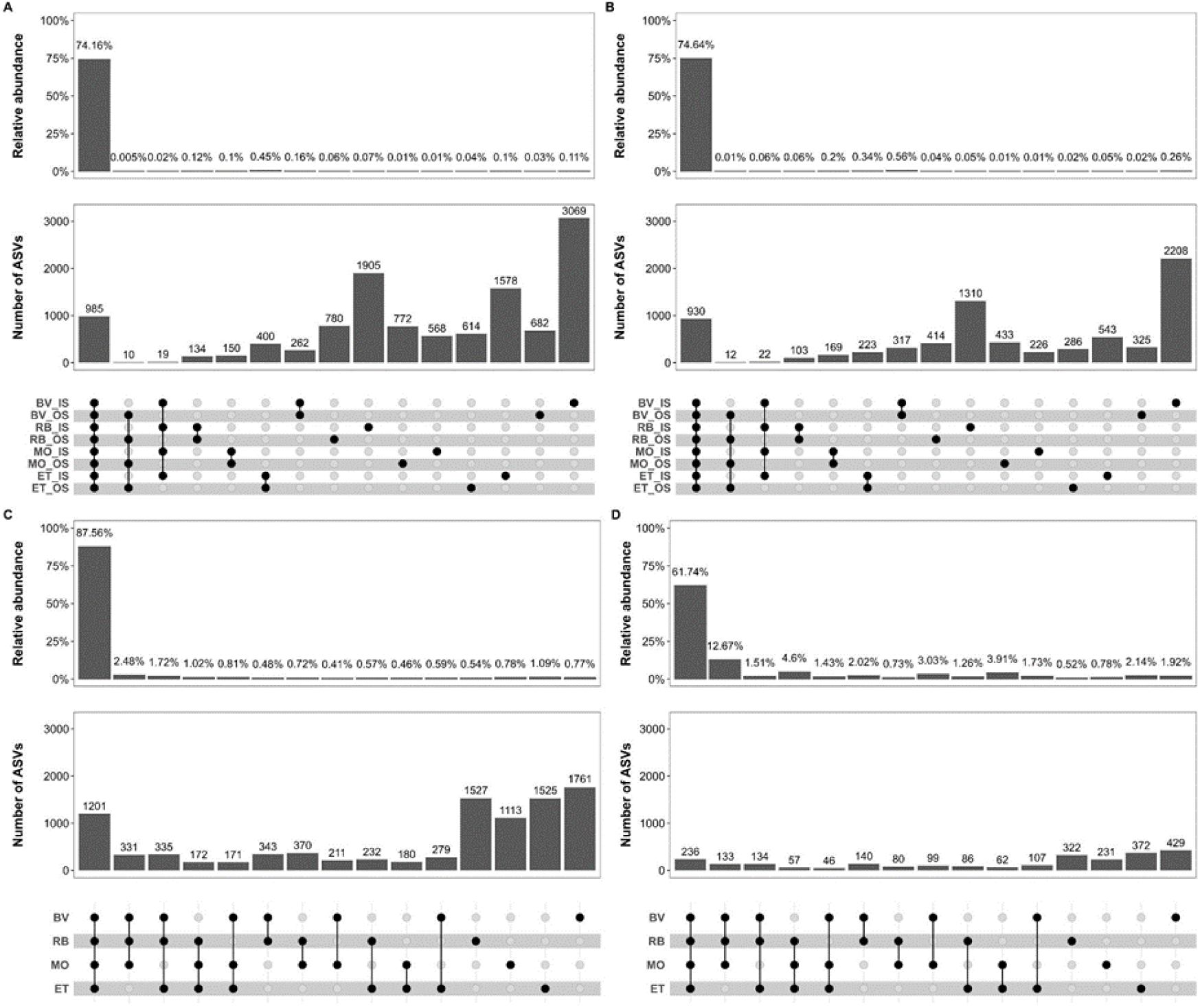
Common and unique ASVs to each sampling condition in waters and oyster tissues. Panels show shared ASVs between sampling sites: (A) bacteria in IS and OS samples, (B) protists in IS and OS samples, (C) bacteria in OY samples, and (D) protists in OY samples. The matrix layout at the bottom indicates specific intersections between sample types (IS and OS) or sites (BV, RB, MO, and ET), with connected dots representing each intersection set. On the X-axis, each bar corresponds to a unique intersection set. The barplot in the lower panel shows the number of shared ASVs within each intersection set. The barplot in the upper panel displays the relative sequence abundance of shared ASVs within each intersection set.

**Supplementary Figure 4.**
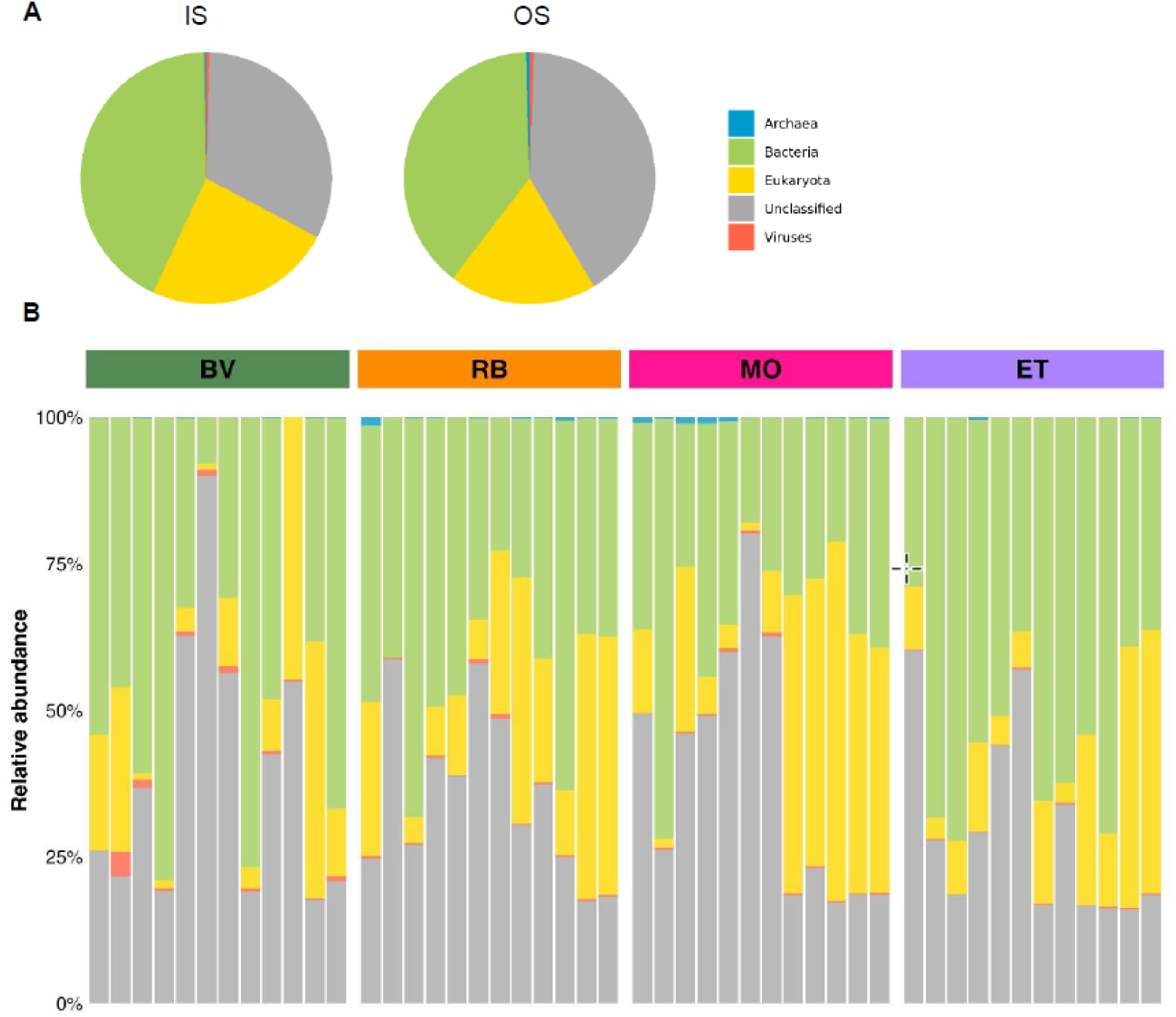
Sequence reads distribution from the metagenomic approach organized by taxonomic domain for water samples categorized, by water type (A) and by sampling site (B). IS, Inshore; OS, Offshore. BV, Bay of Veys; RB, Bay of Brest; MO, Marennes-Oléron; ET, Thau lagoon.

**Supplementary Figure 5.**
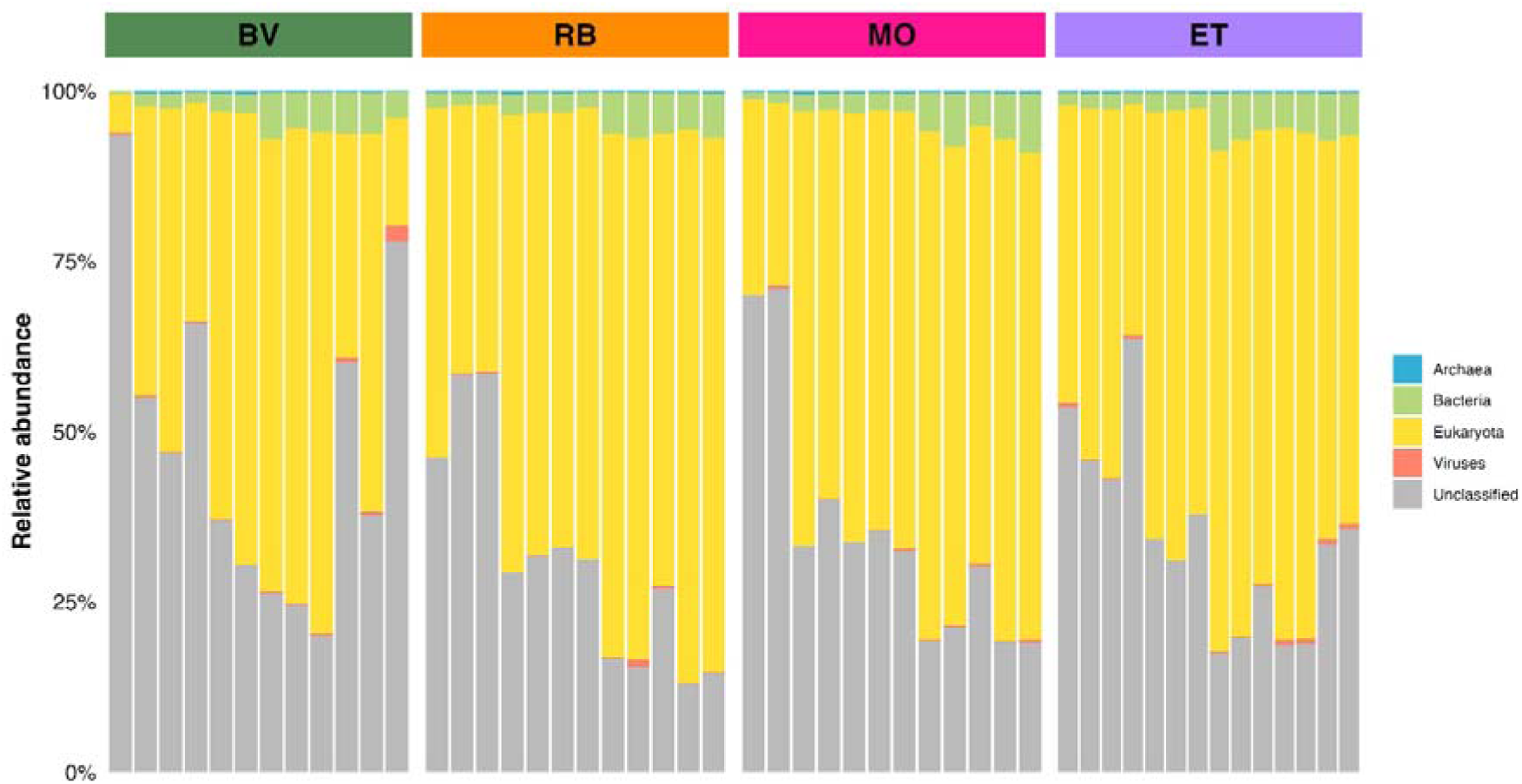
Sequence reads distribution from the metagenomic approach organized by taxonomic domain for oyster samples categorized by site. BV, Bay of Veys; RB, Bay of Brest; MO, Marennes-Oléron; ET, Thau lagoon.

## Notes

https://rome.ifremer.fr/

https://doi.org/10.12770/a14dedb4-65e8-48dc-a07b-be48d546a5df

### Competing Interest Statement

The authors have declared no competing interest.

## List of References

Afssa, Afsset. Évaluation des risques liés à la présence de cyanobactéries et leurs toxines dans les eaux destinées à l’alimentation, à la baignade et autres activités récréatives,. 2006.

Agreste. France Mémento 2023 2024.

Agreste. Enquête Aquaculture 2020. Chiffres et Données. 2021.

Agreste. Nouvelle Aquitaine, Filière Conchyliculture 2018.

Alberti A, Poulain J, Engelen S, Labadie K, Romac S, Ferrera I, et al. Viral to metazoan marine plankton nucleotide sequences from the Tara Oceans expedition. Sci Data 2017;4:170093. 10.1038/sdata.2017.93.

Amadei Martínez L, Sabbe K, D’hondt S, Dasseville R, Daveloose I, Verstraete T, et al. Freshwater Discharge and Salinity Drive Taxonomic and Functional Turnover of Microbial Communities in a Turbid Macrotidal Estuary. Environ Microbiol Rep 2025;17:e70135. 10.1111/1758-2229.70135.

Amzil Z, Derrien A, Terre Terrillon A, Savar V, Bertin T, Peyrat M, et al. Five Years Monitoring the Emergence of Unregulated Toxins in Shellfish in France (EMERGTOX 2018–2022). Mar Drugs 2023;21:435. 10.3390/md21080435.

ANSES. Avis et rapport de l’Anses relatif à l’actualisation de l’évaluation des risques liés aux cyanobactéries et leurs toxines dans les dans les eaux destinées à l’alimentation, les eaux de loisirs et les eaux destinées aux activités de pêche professionnelle et de loisir,. 2020.

Arzul I, Garcia C, Chollet B, Serpin D, Lupo C, Noyer M, et al. First characterization of the parasite *Haplosporidium costale* in France and development of a real-time PCR assay for its rapid detection in the Pacific oyster, *Crassostrea gigas*. Transbound Emerg Dis 2022;69. 10.1111/tbed.14541.

Austin B. Bacterial Pathogens of Marine Fish. In: Belkin S, Colwell RR, editors. Oceans Health Pathog. Mar. Environ., Boston, MA: Springer US; 2005, p. 391–413. 10.1007/0-387-23709-7_17.

Austin B, Austin DA. Bacterial fish pathogens: diseases of farmed and wild fish. 4th ed. Dordrecht: Springer; 2007.

Aylagas E, Borja Á, Irigoien X, Rodríguez-Ezpeleta N. Benchmarking DNA Metabarcoding for Biodiversity-Based Monitoring and Assessment. Front Mar Sci 2016;3. 10.3389/fmars.2016.00096.

Aylagas E, Borja Á, Rodríguez-Ezpeleta N. Environmental Status Assessment Using DNA Metabarcoding: Towards a Genetics Based Marine Biotic Index (gAMBI). PLoS ONE 2014;9:e90529. 10.1371/journal.pone.0090529.

Berry O, Jarman S, Bissett A, Hope M, Paeper C, Bessey C, et al. Making environmental DNA (eDNA) biodiversity records globally accessible. Environ DNA 2021;3:699–705. 10.1002/edn3.173.

Bilbao J, Pavloudi C, Blanco-Rayón E, Franco J, Madariaga I, Seoane S. Phytoplankton community composition in relation to environmental variability in the Urdaibai estuary (SE Bay of Biscay): Microscopy and eDNA metabarcoding. Mar Environ Res 2023;191:106175. 10.1016/j.marenvres.2023.106175.

Biré R, Bertin T, Dom I, Hort V, Schmitt C, Diogène J, et al. First Evidence of the Presence of Anatoxin-A in Sea Figs Associated with Human Food Poisonings in France. Mar Drugs 2020;18:285. 10.3390/md18060285.

Bolyen E, Rideout JR, Dillon MR, Bokulich NA, Abnet CC, Al-Ghalith GA, et al. Reproducible, interactive, scalable and extensible microbiome data science using QIIME 2. Nat Biotechnol 2019;37:852–7. 10.1038/s41587-019-0209-9.

Bonny P, Schaeffer J, Besnard A, Desdouits M, Ngang JJE, Le Guyader FS. Human and Animal RNA Virus Diversity Detected by Metagenomics in Cameroonian Clams. Front Microbiol 2021;12:770385. 10.3389/fmicb.2021.770385.

Bormans M, Amzil Z, Mineaud E, Brient L, Savar V, Robert E, et al. Demonstrated transfer of cyanobacteria and cyanotoxins along a freshwater-marine continuum in France. Harmful Algae 2019;87:101639. 10.1016/j.hal.2019.101639.

Bormans M, Savar V, Legrand B, Mineaud E, Robert E, Lance E, et al. Cyanobacteria and cyanotoxins in estuarine water and sediment. Aquat Ecol 2020;54:625–40. 10.1007/s10452-020-09764-y.

Bracken LJ, Wainwright J, Ali GA, Tetzlaff D, Smith MW, Reaney SM, et al. Concepts of hydrological connectivity: Research approaches, pathways and future agendas. Earth-Sci Rev 2013;119:17–34. 10.1016/j.earscirev.2013.02.001.

Brenner DJ, Boone DR, Garrity GM, Goodfellow M, Krieg NR, Rainey FA, et al., editors. Bergey’s Manual of Systematic Bacteriology: Volume Two The Proteobacteria Part B The Gammaproteobacteria. Second Edition. Boston, MA: Springer US; 2005. 10.1007/0-387-28022-7.

Burke LM, editor. Pilot analysis of global ecosystems: coastal ecosystems. Washington, DC: World Resources Institute; 2001.

Buttigieg PL, Fadeev E, Bienhold C, Hehemann L, Offre P, Boetius A. Marine microbes in 4D — using time series observation to assess the dynamics of the ocean microbiome and its links to ocean health. Curr Opin Microbiol 2018;43:169–85. 10.1016/j.mib.2018.01.015.

Callahan BJ, McMurdie PJ, Rosen MJ, Han AW, Johnson AJA, Holmes SP. DADA2: High-resolution sample inference from Illumina amplicon data. Nat Methods 2016;13:581–3. 10.1038/nmeth.3869.

Campbell BJ, Kirchman DL. Bacterial diversity, community structure and potential growth rates along an estuarine salinity gradient. ISME J 2013;7:210–20. 10.1038/ismej.2012.93.

Caracciolo M, Rigaut-Jalabert F, Romac S, Mahé F, Forsans S, Gac J, et al. Seasonal dynamics of marine protist communities in tidally mixed coastal waters. Mol Ecol 2022;31:3761–83. 10.1111/mec.16539.

Clerissi C, Brunet S, Vidal-Dupiol J, Adjeroud M, Lepage P, Guillou L, et al. Protists Within Corals: The Hidden Diversity. Front Microbiol 2018;9:2043. 10.3389/fmicb.2018.02043.

Cocciardi JM, Hoffman AM, Alvarado-Serrano DF, Anderson J, Blumstein M, Boehm EL, et al. The value of long-term ecological research for evolutionary insights. Nat Ecol Evol 2024;8:1584–92. 10.1038/s41559-024-02464-y.

Cordier T, Alonso-Sáez L, Apothéloz-Perret-Gentil L, Aylagas E, Bohan DA, Bouchez A, et al. Ecosystems monitoring powered by environmental genomics: A review of current strategies with an implementation roadmap. Mol Ecol 2021;30:2937–58. 10.1111/mec.15472.

Cottrell MT, Ras J, Kirchman DL. Bacteriochlorophyll and community structure of aerobic anoxygenic phototrophic bacteria in a particle-rich estuary. ISME J 2010;4:945–54. 10.1038/ismej.2010.13.

Crump BC, Bowen JL. The Microbial Ecology of Estuarine Ecosystems. Annu Rev Mar Sci 2024;16:335–60. 10.1146/annurev-marine-022123-101845.

Derolez V, Bec B, Munaron D, Fiandrino A, Pete R, Simier M, et al. Recovery trajectories following the reduction of urban nutrient inputs along the eutrophication gradient in French Mediterranean lagoons. Ocean Coast Manag 2019;171:1–10. 10.1016/j.ocecoaman.2019.01.012.

Derolez V, Malet N, Fiandrino A, Lagarde F, Richard M, Ouisse V, et al. Fifty years of ecological changes: Regime shifts and drivers in a coastal Mediterranean lagoon during oligotrophication. Sci Total Environ 2020a;732. 10.1016/j.scitotenv.2020.139292.

Derolez V, Soudant D, Malet N, Chiantella C, Richard M, Abadie E, et al. Two decades of oligotrophication: Evidence for a phytoplankton community shift in the coastal lagoon of Thau (Mediterranean Sea, France). Estuar Coast Shelf Sci 2020b;241:106810. 10.1016/j.ecss.2020.106810.

Desdouits M, Piquet J-C, Wacrenier C, Le Mennec C, Parnaudeau S, Jousse S, et al. Can shellfish be used to monitor SARS-CoV-2 in the coastal environment? Sci Total Environ 2021;778:146270. 10.1016/j.scitotenv.2021.146270.

Desdouits M, Reynaud Y, Philippe C, Guyader FSL. A Comprehensive Review for the Surveillance of Human Pathogenic Microorganisms in Shellfish. Microorganisms 2023;11:2218. 10.3390/microorganisms11092218.

Dungan CF, Elston RA. Histopathological and ultrastructural characteristics of bacterial destruction of the hinge ligaments of cultured juvenile Pacific oysters, *Crassostrea gigas*. Aquaculture 1988;72:1–14. 10.1016/0044-8486(88)90141-X.

EFSA Panel on Biological Hazards (BIOHAZ), Koutsoumanis K, Allende A, Alvarez-Ordóñez A, Bolton D, Bover-Cid S, et al. Public health aspects of Vibrio spp. related to the consumption of seafood in the EU. EFSA J 2024;22. 10.2903/j.efsa.2024.8896.

Eisenhofer R, Minich JJ, Marotz C, Cooper A, Knight R, Weyrich LS. Contamination in Low Microbial Biomass Microbiome Studies: Issues and Recommendations. Trends Microbiol 2019;27:105–17. 10.1016/j.tim.2018.11.003.

Escobedo-Hinojosa W, Pardo-López L. Analysis of bacterial metagenomes from the Southwestern Gulf of Mexico for pathogens detection. Pathog Dis 2017;75. 10.1093/femspd/ftx058.

European Food Safety Authority (EFSA). Analysis of the European baseline survey of norovirus in oysters. EFSA J 2019;17. 10.2903/j.efsa.2019.5762.

Fagervold SK, Bourgeois S, Pruski AM, Charles F, Kerhervé P, Vétion G, et al. River organic matter shapes microbial communities in the sediment of the Rhône prodelta. ISME J 2014;8:2327–38. 10.1038/ismej.2014.86.

Farkas K, Cooper DM, McDonald JE, Malham SK, De Rougemont A, Jones DL. Seasonal and spatial dynamics of enteric viruses in wastewater and in riverine and estuarine receiving waters. Sci Total Environ 2018;634:1174–83. 10.1016/j.scitotenv.2018.04.038.

Fiandrino A, Martin Y, Got P, Bonnefont JL, Troussellier M. Bacterial contamination of Mediterranean coastal seawater as affected by riverine inputs: simulation approach applied to a shellfish breeding area (Thau lagoon, France). Water Res 2003;37:1711–22. 10.1016/S0043-1354(02)00573-0.

Fiandrino A, Ouisse V, Dumas F, Lagarde F, Pete R, Malet N, et al. Spatial patterns in coastal lagoons related to the hydrodynamics of seawater intrusion. Mar Pollut Bull 2017;119:132–44. 10.1016/j.marpolbul.2017.03.006.

Ficetola GF, Miaud C, Pompanon F, Taberlet P. Species detection using environmental DNA from water samples. Biol Lett 2008;4:423–5. 10.1098/rsbl.2008.0118.

Fortunato CS, Herfort L, Zuber P, Baptista AM, Crump BC. Spatial variability overwhelms seasonal patterns in bacterioplankton communities across a river to ocean gradient. ISME J 2012;6:554–63. 10.1038/ismej.2011.135.

Foster B, Grewal S, Graves O, Hughes FM, Sokolova IM. Copper exposure affects hemocyte apoptosis and *Perkinsus marinus* infection in eastern oysters *Crassostrea virginica* (Gmelin). Fish Shellfish Immunol 2011;31:341–9. 10.1016/j.fsi.2011.05.024.

Fraga M, Churro C, Leão-Martins J, Rudnitskaya A, Botelho MJ. Cyanotoxins on the move - Freshwater origins with marine consequences: A systematic review of global changes and emerging trends. Mar Pollut Bull 2025;216:118017. 10.1016/j.marpolbul.2025.118017.

Garate L, Alonso-Sáez L, Revilla M, Logares R, Lanzén A. Shared and contrasting associations in the dynamic nano- and picoplankton communities of two close but contrasting sites from the Bay of Biscay. Environ Microbiol 2022;24:6052–70. 10.1111/1462-2920.16153.

Gilbert JA, Field D, Swift P, Thomas S, Cummings D, Temperton B, et al. The Taxonomic and Functional Diversity of Microbes at a Temperate Coastal Site: A ‘Multi-Omic’ Study of Seasonal and Diel Temporal Variation. PLoS ONE 2010;5:e15545. 10.1371/journal.pone.0015545.

Gobet A, Böer SI, Huse SM, van Beusekom JEE, Quince C, Sogin ML, et al. Diversity and dynamics of rare and of resident bacterial populations in coastal sands. ISME J 2012;6:542–53. 10.1038/ismej.2011.132.

Gómez-Chiarri M, Andrews JS, Coppersmith J, Guidry ME, Houtz A, Mills B, et al. Vibriosis of bivalves. Dis. Bivalves, Elsevier; 2025, p. 143–62. 10.1016/B978-0-12-820339-2.00005-X.

Granit J, Liss Lymer B, Olsen S, Tengberg A, Nõmmann S, Clausen TJ. A conceptual framework for governing and managing key flows in a source-to-sea continuum. Water Policy 2017;19:673–91. 10.2166/wp.2017.126.

Guillou L, Bachar D, Audic S, Bass D, Berney C, Bittner L, et al. The Protist Ribosomal Reference database (PR2): a catalog of unicellular eukaryote small sub-unit rRNA sequences with curated taxonomy. Nucleic Acids Res 2013;41:D597–604. 10.1093/nar/gks1160.

He Q, Silliman BR. Climate Change, Human Impacts, and Coastal Ecosystems in the Anthropocene. Curr Biol 2019;29:R1021–35. 10.1016/j.cub.2019.08.042.

Herlemann DPR, Labrenz M, Jürgens K, Bertilsson S, Waniek JJ, Andersson AF. Transitions in bacterial communities along the 2000 km salinity gradient of the Baltic Sea. ISME J 2011;5:1571–9. 10.1038/ismej.2011.41.

Hervé V, Morelle J, Lambourdière J, Lopez PJ, Claquin P. Together throughout the year: seasonal patterns of bacterial and eukaryotic microbial communities in a macrotidal estuary. Environ Microbiome 2025;20:8. 10.1186/s40793-025-00664-y.

Ibarbalz FM, Henry N, Brandão MC, Martini S, Busseni G, Byrne H, et al. Global Trends in Marine Plankton Diversity across Kingdoms of Life. Cell 2019;179:1084–1097.e21. 10.1016/j.cell.2019.10.008.

Izquierdo-Lara RW, Worp N, Olthof M, Oude Munnink BB, Schapendonk CME, Prasad DK, et al. Tracking norovirus diversity at a global scale through wastewater metagenomics. Water Res 2025;287:124257. 10.1016/j.watres.2025.124257.

Jiajun L, Biao Z, Guangshuai Z, Sihui S, Yansong L, Jinhui Z, et al. Flooding promotes the coalescence of microbial community in estuarine habitats. Mar Environ Res 2024;202:106735. 10.1016/j.marenvres.2024.106735.

John SG, Mendez CB, Deng L, Poulos B, Kauffman AKM, Kern S, et al. A simple and efficient method for concentration of ocean viruses by chemical flocculation. Environ Microbiol Rep 2011;3:195–202. 10.1111/j.1758-2229.2010.00208.x.

Katayama H, Shimasaki A, Ohgaki S. Development of a Virus Concentration Method and Its Application to Detection of Enterovirus and Norwalk Virus from Coastal Seawater. Appl Environ Microbiol 2002;68:1033–9. 10.1128/AEM.68.3.1033-1039.2002.

King WL, Jenkins C, Seymour JR, Labbate M. Oyster disease in a changing environment: Decrypting the link between pathogen, microbiome and environment. Mar Environ Res 2019;143:124–40. 10.1016/j.marenvres.2018.11.007.

Kirchman D, Cottrel M, DiTullio G. Shaping of bacterial community composition and diversity by phytoplankton and salinity in the Delaware Estuary, USA. Aquat Microb Ecol 2017;78:93–106. 10.3354/ame01805.

Klindworth A, Pruesse E, Schweer T, Peplies J, Quast C, Horn M, et al. Evaluation of general 16S ribosomal RNA gene PCR primers for classical and next-generation sequencing-based diversity studies. Nucleic Acids Res 2013;41:e1–e1. 10.1093/nar/gks808.

La Jeunesse I, Elliott M. Anthropogenic regulation of the phosphorus balance in the Thau catchment-coastal lagoon system (Mediterraean Sea, France) over 24 years. Mar Pollut Bull 2004;48:679–87. 10.1016/j.marpolbul.2003.10.011.

Lassus P, Chomérat N, Hess P, Nézan E. Toxic and harmful microalgae of the world ocean: = Micro-algues toxiques et nuisibles de l’océan mondial. Copenhagen N, Denmark: International Society for the Study of Harmful Algae; 2016.

Latouche C, Jouanneau JM. Synthèse des connaissances de l’estuaire de la Gironde. 1990.

Leight AK, Crump BC, Hood RR. Assessment of Fecal Indicator Bacteria and Potential Pathogen Co-Occurrence at a Shellfish Growing Area. Front Microbiol 2018;9:384. 10.3389/fmicb.2018.00384.

Li Y, Bhatt P, Xagoraraki I. In-depth comparison of untargeted and targeted sequencing for detecting virus diversity in wastewater. Water Res 2025;283:123803. 10.1016/j.watres.2025.123803.

Lima-Mendez G, Faust K, Henry N, Decelle J, Colin S, Carcillo F, et al. Determinants of community structure in the global plankton interactome. Science 2015;348:1262073. 10.1126/science.1262073.

Loch TP, Faisal M. Emerging flavobacterial infections in fish: A review. J Adv Res 2015;6:283–300. 10.1016/j.jare.2014.10.009.

Logares R, Bråte J, Bertilsson S, Clasen JL, Shalchian-Tabrizi K, Rengefors K. Infrequent marine–freshwater transitions in the microbial world. Trends Microbiol 2009;17:414–22. 10.1016/j.tim.2009.05.010.

Logares R, Deutschmann IM, Junger PC, Giner CR, Krabberød AK, Schmidt TSB, et al. Disentangling the mechanisms shaping the surface ocean microbiota. Microbiome 2020;8:55. 10.1186/s40168-020-00827-8.

Luo Z, Krock B, Mertens KN, Nézan E, Chomérat N, Bilien G, et al. Adding new pieces to the *Azadinium* (Dinophyceae) diversity and biogeography puzzle: Non-toxigenic *Azadinium zhuanum* sp. nov. from China, toxigenic *A. poporum* from the Mediterranean, and a non-toxigenic *A. dalianense* from the French Atlantic. Harmful Algae 2017;66:65–78. 10.1016/j.hal.2017.05.001.

Marshall HG, Burchardt L, Lacouture R. A review of phytoplankton composition within Chesapeake Bay and its tidal estuaries. J Plankton Res 2005;27:1083–102. 10.1093/plankt/fbi079.

Martin M. Cutadapt removes adapter sequences from high-throughput sequencing reads. EMBnetJournal 2011;17:10. 10.14806/ej.17.1.200.

Martínez ML, Intralawan A, Vázquez G, Pérez-Maqueo O, Sutton P, Landgrave R. The coasts of our world: Ecological, economic and social importance. Ecol Econ 2007;63:254–72. 10.1016/j.ecolecon.2006.10.022.

Mason OU, Canter EJ, Gillies LE, Paisie TK, Roberts BJ. Mississippi River Plume Enriches Microbial Diversity in the Northern Gulf of Mexico. Front Microbiol 2016;7. 10.3389/fmicb.2016.01048.

Massana R, Gobet A, Audic S, Bass D, Bittner L, Boutte C, et al. Marine protist diversity in European coastal waters and sediments as revealed by high-throughput sequencing. Environ Microbiol 2015;17:n/a-n/a. 10.1111/1462-2920.12955.

Mauffret A, Caprais M-P, Gourmelon M. Relevance of Bacteroidales and F-Specific RNA Bacteriophages for Efficient Fecal Contamination Tracking at the Level of a Catchment in France. Appl Environ Microbiol 2012;78:5143–52. 10.1128/AEM.00315-12.

McMurdie PJ, Holmes S. phyloseq: An R Package for Reproducible Interactive Analysis and Graphics of Microbiome Census Data. PLoS ONE 2013;8:e61217. 10.1371/journal.pone.0061217.

Mérou N, Lecadet C, Ubertini M, Pouvreau S, Arzul I. Environmental distribution and seasonal dynamics of Marteilia refringens and Bonamia ostreae, two protozoan parasites of the European flat oyster, Ostrea edulis. Front Cell Infect Microbiol 2023;13:1154484. 10.3389/fcimb.2023.1154484.

Meyneng M, Lemonnier H, Gendre RL, Plougoulen G, Antypas F, Ansquer D, et al. Subtropical coastal microbiome variations due to massive river runoff after a cyclonic event. Environ Microbiome 2024;19:10. 10.1186/s40793-024-00554-9.

Moreau P, Moreau K, Segarra A, Tourbiez D, Travers M-A, Rubinsztein DC, et al. Autophagy plays an important role in protecting Pacific oysters from OsHV-1 and *Vibrio aestuarianus* infections. Autophagy 2015;11:516–26. 10.1080/15548627.2015.1017188.

Nézan E, Siano R, Boulben S, Six C, Bilien G, Chèze K, et al. Genetic diversity of the harmful family Kareniaceae (Gymnodiniales, Dinophyceae) in France, with the description of *Karlodinium gentienii* sp. nov.: A new potentially toxic dinoflagellate. Harmful Algae 2014;40:75–91. 10.1016/j.hal.2014.10.006.

Nieuwenhuijse DF, Oude Munnink BB, Phan MVT, the Global Sewage Surveillance project consortium, Hendriksen RS, Bego A, et al. Setting a baseline for global urban virome surveillance in sewage. Sci Rep 2020;10:13748. 10.1038/s41598-020-69869-0.

Nogales B, Lanfranconi MP, Piña-Villalonga JM, Bosch R. Anthropogenic perturbations in marine microbial communities. FEMS Microbiol Rev 2011;35:275–98. 10.1111/j.1574-6976.2010.00248.x.

Olesen JM, Bascompte J, Dupont YL, Jordano P. The modularity of pollination networks. Proc Natl Acad Sci 2007;104:19891–6. 10.1073/pnas.0706375104.

Paillard C, Gueguen Y, Wegner KM, Bass D, Pallavicini A, Vezzulli L, et al. Recent advances in bivalve-microbiota interactions for disease prevention in aquaculture. Curr Opin Biotechnol 2022;73:225–32. 10.1016/j.copbio.2021.07.026.

Paillard C, Le Roux F, Borrego JJ. Bacterial disease in marine bivalves, a review of recent studies: Trends and evolution. Aquat Living Resour 2004;17:477–98. 10.1051/alr:2004054.

Parada AE, Needham DM, Fuhrman JA. Every base matters: assessing small subunit rRNA primers for marine microbiomes with mock communities, time series and global field samples: Primers for marine microbiome studies. Environ Microbiol 2016;18:1403–14. 10.1111/1462-2920.13023.

Pawlowski J, Apothéloz-Perret-Gentil L, Mächler E, Altermatt F. Environmental DNA applications for biomonitoring and bioassessment in aquatic ecosystems 2020. 10.5167/UZH-187800.

Pawlowski J, Bonin A, Boyer F, Cordier T, Taberlet P. Environmental DNA for biomonitoring. Mol Ecol 2021;30:2931–6. 10.1111/mec.16023.

Pellouin-Grouhel A, Romana A. Mise en œuvre de la DCE dans les zones littorales□: préconisations pour le contrôle de surveillance et éléments pour le contrôle opérationnel. Houille Blanche 2006;92:68–74. 10.1051/lhb:200604011.

Pete R, Guyondet T, Bec B, Derolez V, Cesmat L, Lagarde F, et al. A box-model of carrying capacity of the Thau lagoon in the context of ecological status regulations and sustainable shellfish cultures. Ecol Model 2020;426:109049–109049. 10.1016/j.ecolmodel.2020.109049.

Plus M, Jeunesse IL, Bouraoui F, Zaldívar JM, Chapelle A, Lazure P. Modelling water discharges and nitrogen inputs into a Mediterranean lagoon: Impact on the primary production. Ecol Model 2006;193:69–89. 10.1016/j.ecolmodel.2005.07.037.

Preece EP, Hardy FJ, Moore BC, Bryan M. A review of microcystin detections in Estuarine and Marine waters: Environmental implications and human health risk. Harmful Algae 2017;61:31–45. 10.1016/j.hal.2016.11.006.

Priest T, Henry N, Weber T, Planat L, Rousseau C, Dittami SM, et al. The JEDI marker as a universal measure of planetary biodiversity 2025. 10.1101/2025.08.11.669668.

Pritchard DW. Estuarine Classification — A Help or a Hindrance. In: Neilson BJ, Kuo A, Brubaker J, editors. Estuar. Circ., Totowa, NJ: Humana Press; 1989, p. 1–38. 10.1007/978-1-4612-4562-9_1.

Pruesse E, Quast C, Knittel K, Fuchs BM, Ludwig W, Peplies J, et al. SILVA: a comprehensive online resource for quality checked and aligned ribosomal RNA sequence data compatible with ARB. Nucleic Acids Res 2007;35:7188–96. 10.1093/nar/gkm864.

Ramond P, Sourisseau M, Simon N, Romac S, Schmitt S, Rigaut-Jalabert F, et al. Coupling between taxonomic and functional diversity in protistan coastal communities: Functional diversity of marine protists. Environ Microbiol 2019;21:730–49. 10.1111/1462-2920.14537.

Ratiskol G. Suivi des efflorescences phytoplanctoniques dans le bassin de Marennes-Oléron en 1989. 1994.

Reignier O, Bormans M, Hervé F, Robert E, Savar V, Tanniou S, et al. Spatio-temporal connectivity of a toxic cyanobacterial community and its associated microbiome along a freshwater-marine continuum. Harmful Algae 2024;134:102627. 10.1016/j.hal.2024.102627.

Rincé A, Balière C, Hervio-Heath D, Cozien J, Lozach S, Parnaudeau S, et al. Occurrence of Bacterial Pathogens and Human Noroviruses in Shellfish-Harvesting Areas and Their Catchments in France. Front Microbiol 2018;9:2443. 10.3389/fmicb.2018.02443.

Ríos-Castro R, Cabo A, Teira E, Cameselle C, Gouveia S, Payo P, et al. High-throughput sequencing as a tool for monitoring prokaryote communities in a wastewater treatment plant. Sci Total Environ 2023;861:160531. 10.1016/j.scitotenv.2022.160531.

Ryckman M, Gantois N, Dominguez RG, Desramaut J, Li L-L, Even G, et al. Molecular Identification and Subtype Analysis of Blastocystis sp. Isolates from Wild Mussels (Mytilus edulis) in Northern France. Microorganisms 2024;12:710. 10.3390/microorganisms12040710.

Salter SJ, Cox MJ, Turek EM, Calus ST, Cookson WO, Moffatt MF, et al. Reagent and laboratory contamination can critically impact sequence-based microbiome analyses. BMC Biol 2014;12:87. 10.1186/s12915-014-0087-z.

Savio D, Stadler P, Reischer GH, Demeter K, Linke RB, Blaschke AP, et al. Spring Water of an Alpine Karst Aquifer Is Dominated by a Taxonomically Stable but Discharge-Responsive Bacterial Community. Front Microbiol 2019;10:28. 10.3389/fmicb.2019.00028.

Schaeffer J, Desdouits M, Besnard A, Le Guyader FS. Looking into sewage: how far can metagenomics help to detect human enteric viruses? Front Microbiol 2023;14. 10.3389/fmicb.2023.1161674.

Sharp JH, Yoshiyama K, Parker AE, Schwartz MC, Curless SE, Beauregard AY, et al. A Biogeochemical View of Estuarine Eutrophication: Seasonal and Spatial Trends and Correlations in the Delaware Estuary. Estuaries Coasts 2009;32:1023–43. 10.1007/s12237-009-9210-8.

Simon M, Jardillier L, Deschamps P, Moreira D, Restoux G, Bertolino P, et al. Complex communities of small protists and unexpected occurrence of typical marine lineages in shallow freshwater systems. Environ Microbiol 2015;17:3610–27. 10.1111/1462-2920.12591.

Smith MW, Herfort L, Tyrol K, Suciu D, Campbell V, Crump BC, et al. Seasonal Changes in Bacterial and Archaeal Gene Expression Patterns across Salinity Gradients in the Columbia River Coastal Margin. PLoS ONE 2010;5:e13312. 10.1371/journal.pone.0013312.

Sogin ML, Morrison HG, Huber JA, Mark Welch D, Huse SM, Neal PR, et al. Microbial diversity in the deep sea and the underexplored “rare biosphere”. Proc Natl Acad Sci U S A 2006;103:12115–20. 10.1073/pnas.0605127103.

Souchu P, Gasc A, Collos Y, Vaquer A, Tournier H, Bibent B, et al. Biogeochemical aspects of bottom anoxia in a Mediterranean lagoon (Thau, France). Mar Ecol Prog Ser 1998;164:135–46. 10.3354/meps164135.

Stoeck T, Bass D, Nebel M, Christen R, Jones MDM, Breiner H-W, et al. Multiple marker parallel tag environmental DNA sequencing reveals a highly complex eukaryotic community in marine anoxic water. Mol Ecol 2010;19:21–31. 10.1111/j.1365-294X.2009.04480.x.

Strittmatter M. Simultanuous DNA and RNA extraction from various tissues of the two oyster species, Crassostrea gigas and Ostrea edulis, for metabarcoding v1 2019. 10.17504/protocols.io.22aggae.

Strubbia S, Phan MVT, Schaeffer J, Koopmans M, Cotten M, Le Guyader FS. Characterization of Norovirus and Other Human Enteric Viruses in Sewage and Stool Samples Through Next-Generation Sequencing. Food Environ Virol 2019. 10.1007/s12560-019-09402-3.

Strubbia S, Schaeffer J, Besnard A, Wacrenier C, Le Mennec C, Garry P, et al. Metagenomic to evaluate norovirus genomic diversity in oysters: Impact on hexamer selection and targeted capture-based enrichment. Int J Food Microbiol 2020;323:108588. 10.1016/j.ijfoodmicro.2020.108588.

Taberlet P, Bonin A, Zinger L, Coissac E. Environmental DNA: For Biodiversity Research and Monitoring. 1st ed. Oxford University PressOxford; 2018. 10.1093/oso/9780198767220.001.0001.

Takano T, Matsuyama T, Sakai T, Nakamura Y, Kamaishi T, Nakayasu C, et al. Ichthyobacterium seriolicida gen. nov., sp. nov., a member of the phylum ‘Bacteroidetes’, isolated from yellowtail fish (Seriola quinqueradiata) affected by bacterial haemolytic jaundice, and proposal of a new family, Ichthyobacteriaceae fam. nov. Int J Syst Evol Microbiol 2016;66:580–6. 10.1099/ijsem.0.000757.

Tee HS, Waite D, Lear G, Handley KM. Microbial river-to-sea continuum: gradients in benthic and planktonic diversity, osmoregulation and nutrient cycling. Microbiome 2021;9:190. 10.1186/s40168-021-01145-3.

Telesh IV, Khlebovich VV. Principal processes within the estuarine salinity gradient: A review. Mar Pollut Bull 2010;61:149–55. 10.1016/j.marpolbul.2010.02.008.

They NH, Ferreira LMH, Marins LF, Abreu PC. Bacterial Community Composition and Physiological Shifts Associated with the El Niño Southern Oscillation (ENSO) in the Patos Lagoon Estuary. Microb Ecol 2015;69:525–34. 10.1007/s00248-014-0511-5.

Tisza M, Javornik Cregeen S, Avadhanula V, Zhang P, Ayvaz T, Feliz K, et al. Wastewater sequencing reveals community and variant dynamics of the collective human virome. Nat Commun 2023;14:6878. 10.1038/s41467-023-42064-1.

Travers M-A, Boettcher Miller K, Roque A, Friedman CS. Bacterial diseases in marine bivalves. J Invertebr Pathol 2015;131:11–31. 10.1016/j.jip.2015.07.010.

Urvoy M, Gourmelon M, Serghine J, Rabiller E, L’Helguen S, Labry C. Free-living and particle-attached bacterial community composition, assembly processes and determinants across spatiotemporal scales in a macrotidal temperate estuary. Sci Rep 2022;12:13897. 10.1038/s41598-022-18274-w.

de Vargas C, Audic S, Henry N, Decelle J, Mahé F, Logares R, et al. Eukaryotic plankton diversity in the sunlit ocean. Science 2015;348:1261605. 10.1126/science.1261605.

Vela AI, Fernández A, Bernaldo De Quirós Y, Herráez P, Domínguez L, Fernández-Garayzábal JF. Weissella ceti sp. nov., isolated from beaked whales (Mesoplodon bidens). Int J Syst Evol Microbiol 2011;61:2758–62. 10.1099/ijs.0.028522-0.

Verdier H, Konecny-Dupre L, Marquette C, Reveron H, Tadier S, Grémillard L, et al. Passive sampling of environmental DNA in aquatic environments using 3D-printed hydroxyapatite samplers. Mol Ecol Resour 2022;22:2158–70. 10.1111/1755-0998.13604.

Walker JR, Bente DA, Burch MT, Cerqueira FM, Ren P, Labonté JM. Molecular assessment of oyster microbiomes and viromes reveals their potential as pathogen and ecological sentinels. One Health 2025;20:100973. 10.1016/j.onehlt.2025.100973.

Wang H, Chen F, Zhang C, Wang M, Kan J. Estuarine gradients dictate spatiotemporal variations of microbiome networks in the Chesapeake Bay. Environ Microbiome 2021;16:22. 10.1186/s40793-021-00392-z.

Wegner KM, Morga B, Guillou L, Strittmatter M, Lecadet C, Travers M-A, et al. Prokaryotic microbiota outperform eukaryotic microbiota in differentiating between infection states of iconic diseases of two commercial oyster species. Aquaculture 2025;594:741363. 10.1016/j.aquaculture.2024.741363.

Whitman WB. Bergey’s Manual of Systematic Bacteriology: Volume 5: the Actinobacteria. 2nd ed. New York, NY: Springer New York; 2012.

Wickham H. ggplot2: elegant graphics for data analysis. Second edition. Cham: Springer international publishing; 2016.

Wildlife Conservation Society. One World, One Health: The Manhattan Principles. One World One Health. 2004.

Wu Z, Li M, Qu L, Zhang C, Xie W. Metagenomic insights into microbial adaptation to the salinity gradient of a typical short residence-time estuary. Microbiome 2024;12:115. 10.1186/s40168-024-01817-w.

Yousaf M, Wang J, Rehman A, Li Z. Microcystins in transitional and marine ecosystems: Source categories, distribution patterns, and ecological impacts. Mar Pollut Bull 2025;220:118432. 10.1016/j.marpolbul.2025.118432.

Zinger L, Gobet A, Pommier T. Two decades of describing the unseen majority of aquatic microbial diversity: SEQUENCING AQUATIC MICROBIAL DIVERSITY. Mol Ecol 2012;21:1878–96. 10.1111/j.1365-294X.2011.05362.x.

Zoppini A, Ademollo N, Bensi M, Berto D, Bongiorni L, Campanelli A, et al. Impact of a river flood on marine water quality and planktonic microbial communities. Estuar Coast Shelf Sci 2019;224:62–72. 10.1016/j.ecss.2019.04.038.

